# Substrate specificity and protein stability drive the divergence of plant-specific DNA methyltransferases

**DOI:** 10.1101/2024.07.11.603080

**Authors:** Jianjun Jiang, Jia Gwee, Jian Fang, Sarah M. Leichter, Dean Sanders, Xinrui Ji, Jikui Song, Xuehua Zhong

**Author notes:** These authors contributed equally (Jianjun Jiang, Jia Gwee). Correspondence (X.Z.), (J.S.).

## Abstract

DNA methylation is an important epigenetic mechanism essential for transposon silencing and genome integrity. Across evolution, the substrates of DNA methylation have diversified between kingdoms to account for genome complexity. In plants, Chromomethylase3 (CMT3) and CMT2 are the major methyltransferases mediating CHG and CHH methylation, respectively. However, how these two enzymes diverge on substrate specificities during evolution remains unknown. Here, we reveal that CMT2 originates from a duplication of the evolutionarily more ancient CMT3 in flowering plants. Lacking a key arginine residue recognizing CHG in CMT2 impairs its CHG methylation activity in most flowering plants. An engineered V1200R mutation empowers CMT2 to restore both CHG and CHH methylation in *Arabidopsis cmt2cmt3* mutant, testifying a loss-of-function effect for CMT2 after ∼200 million years of evolution. Interestingly, CMT2 has evolved a long and unstructured N-terminus critical for balancing protein stability, especially under heat stress. Furthermore, CMT2 N-terminus is plastic and can be tolerant to various natural mutations. Together, this study reveals the mechanism of chromomethylase divergence for context-specific DNA methylation in plants and sheds important lights on DNA methylation evolution and function.

## Introduction

DNA methylation is an important gene regulatory mechanism and plays critical roles in many biological processes such as development, transposon silencing, and genome integrity ^1,2^. Dysregulation of DNA methylation can lead to pleiotropic developmental defects in plants and the development of diseases such as cancer in mammals ^3,4^. While most methylated DNA in mammals is found in the CG context, DNA methylation in plants occurs in CG, CHG, and CHH (H=A,T,C). In *Arabidopsis thaliana*, this complex nature of substrate preference is facilitated by multiple DNA methyltransferases including DOMAINS REARRANGED METHYLTRANSFERASE 2 (DRM2), METHYLTRANSFERASE 1 (MET1), CHROMOMETHYLASE 3 (CMT3), and CMT2 ^5,6^.

Chromomethylases are plant-specific DNA methyltransferases containing a chromo domain, a bromo-adjacent homology (BAH) domain, and a catalytic methyltransferase domain^7,8^. In *A. thaliana*, there are three CMT genes, *CMT1*, *CMT2*, and *CMT3* that emerged from duplication events through the evolution of green plants ^9^. Genome duplication events such as whole genome duplication and small-scale duplication are abundant in plants and are thought to be a driving force for diversity and speciation ^10^. While most duplicated genes become silenced or pseudogenes, functional diversification such as subfunctionalization and neofunctionalization serves as potential mechanism behind duplicate retention ^11^. In the case of *A. thaliana*, the *CMT1* gene is dispensable for DNA methylation in *A. thaliana*, while CMT2 and CMT3 appear to have diversified for the labor division for non-CG methylation ^12-14^. However, how the two CMTs have diversified to confer the increasing complexity of non-CG methylation during plant evolution remains unknown.

Functionally, CMT3 maintains symmetric CHG methylation on transposable elements (TEs) and repetitive sequences in the heterochromatin ^13,14^. Recent studies have also shown that CMT3 is involved in the *de novo* establishment of gene-body methylation ^9,15,16^. On the other hand, CMT2 maintains the asymmetric CHH methylation alongside DRM2, with CMT2 methylating DNA within long TEs in heterochromatic regions whereas DRM2 mediates methylation within short TEs and at the edges of long TEs ^13^. CHH methylation plays important roles in TE silencing and environmental adaptation, and both CMT2 and DRM2 pathways enable a “double-lock mechanism”, indicated by the conversion of CMT2 targets to DRM2 targets during the loss of remodeler DDM1, for ensuring maintenance of CHH methylation and genome integrity ^17-19^. Interestingly, CMT2 is also capable of maintaining CHG methylation alongside CMT3, suggesting a partial redundancy between the two chromomethylases to ensure maintenance of methylation in the heterochromatin for genome stability ^14^.

Although CMT2 and CMT3 have distinct DNA substrate preferences, both proteins recognize and bind to methylated histone 3 lysine 9 (H3K9me) through both chromo and BAH domains to methylate DNA substrate ^12,14,20^. The substrate specificity of CMT3 has recently been illustrated in maize homolog, ZMET2, where the enzyme is activated by allosteric recognition of H3K9me2 and histone 3 lysine 18 (H3K18), and the base-specific interactions of the methyltransferase domain with the hemi-methylated CHG site and deformation of the DNA around the target cytosine for methylation ^21^.

Interestingly, the presence of CMT2 and CMT3 differs between plant species; CMT3 is present in green plant lineages ranging from algae to flowering plants (angiosperms), while CMT2 is only present in angiosperms ^9,22,23^, suggesting that CMT3 is more ancient and that CMT2 may arise from CMT3 duplication during evolution. However, not all angiosperm species possess both CMT2 and CMT3; *Zea mays* lost CMT2 even though its close relative *Sorghum bicolor* retains it, while two close Brassicaceae relative of *A. thaliana*, *Eutrema salsugineum* and *Conringia planisiliqua*, lost their CMT3 ^9,13,16^. Furthermore, some angiosperm species such as *Oryza sativa* contains multiple copies of CMT3 ^24^, yet the basis and consequence behind the variation and retention of CMTs in various angiosperm species remain unknown.

Here, we investigated the molecular mechanism underlying the divergence of CMT2 and CMT3 by carrying out a comprehensive structural, functional, and evolutionary study. We noted an arginine residue crucial for the recognition of CHG by CMT3 (R745) showed great variations in CMT2, explaining its loss of CHG specificity. Mutation of the corresponding residue in CMT2 to arginine in *Arabidopsis* (V1200R) gained CHG methylation activity and re-silenced a subset of TEs in *cmt2 cmt3* mutant with CMT3-like function. While CMT3 has a short N-terminus, CMT2 contains a long and disordered N-terminus, which is a common characteristic among many plant species. This long N-terminus regulated CMT2 stability and mediated heat-induced CMT2 degradation. Furthermore, CMT2 N-terminus is more plastic and tolerant to mutations as various CMT2 variations at the N-terminus are observed in nature. Together, this study reveals the mechanism of chromomethylase divergence and provides important insights into DNA methylation function and evolution in plants.

## Results

### CMT2 is evolved and duplicated from CMT3

Our recent structural and functional study of maize CMT3 (ZMET2) homolog has revealed critical residues responsible for its enzymatic preference for CHG substrates ^21^. Using these key residues as a signature, we performed a phylogenetic analysis of CMT3 and CMT2 from a range of species representing major plant lineages across the evolution (Supplementary Dataset 1). We found that CMT3 is present in all green plants (Viridiplantae), including chlorophytes and streptophytes, whereas CMT2 only appears in flowering plants (Figure 1A and Supplementary Dataset 1), consistent with previous reports ^9,23^. Key residues of Y776, R804, and H808 for CHG substrate recognition and W224, Y302, and M641 for H3 binding in ZMET2, are highly conserved across streptophytes, including charophytes and land plants, but undergo notable variations in chlorophyte green algae (Supplemental Figure 1A), suggesting an early emergence of CMT3 as a conserved DNA methyltransferase in green plants. All but one of these ZMET2 residues are also conserved in angiosperm CMT2 (Supplemental Figure 1B), further supporting the existence of CMT3 before CMT2. To examine whether CMT3 homologs from the ancient plant species are functional, we ectopically expressed *UBQ10* promoter-driven coding sequences of CMT3 from a charophyte species (*Chara braunii,* CbCMT3) and a bryophyte species (*Marchantia polymorpha,* MpCMT3) and transformed them into *Arabidopsis cmt3* mutant (Supplementary Figure 2A,B). We next performed an McrBC (an enzyme that specifically cleaves methylated DNA) digestion assay and found that MpCMT3 was able to partially methylate DNA on *Cluster4* (a well-known CMT3 target site), while CbCMT3 showed little methylation activity (Supplementary Figure 2C). These results suggest that additional layers of regulation other than the examined conserved residues are involved in the methylation activity during evolution.

**Figure 1.**
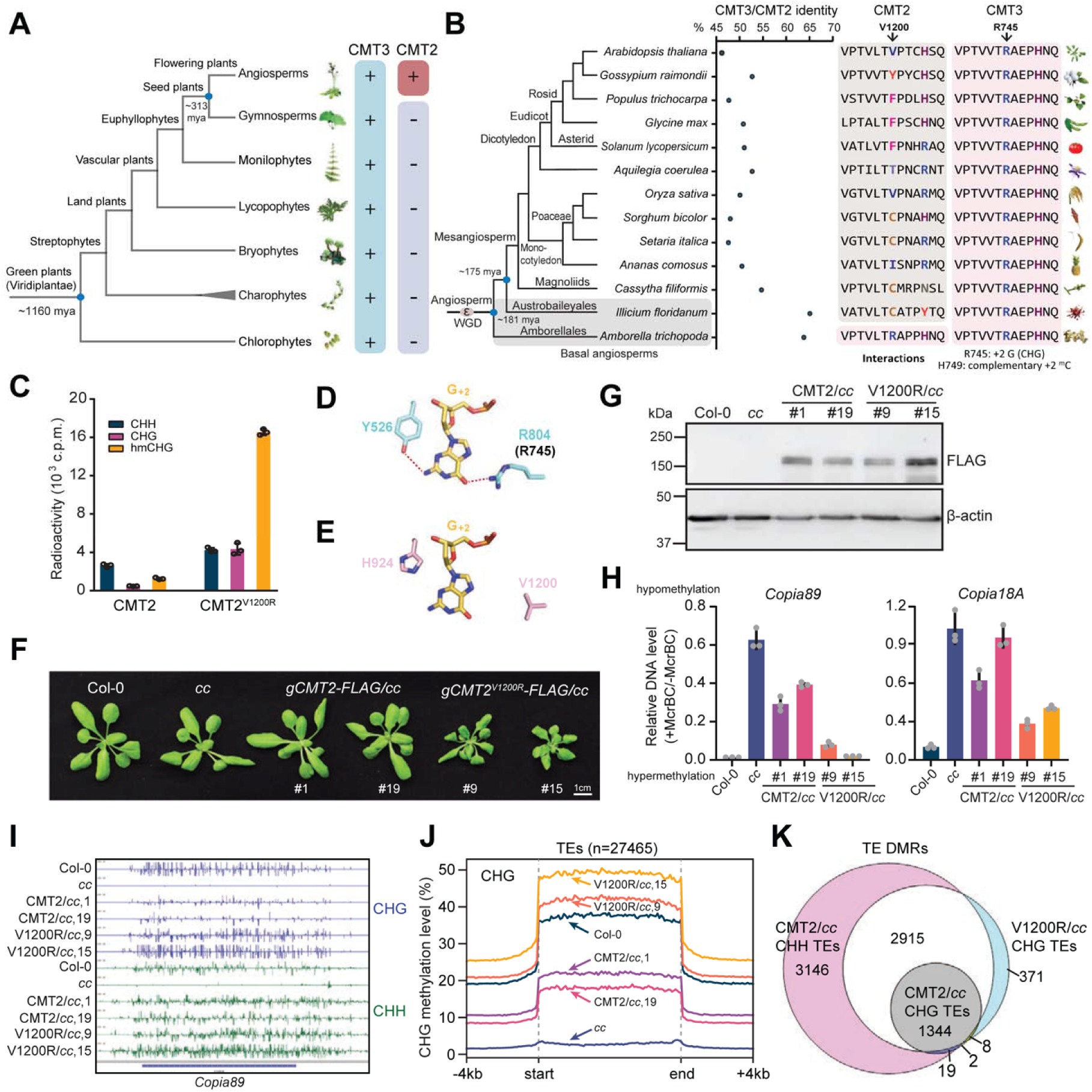
Loss of an arginine residue confers CMT2 a CMT3-distinct substrate preference. (A) A phylogenetic tree depicting the presence of CMT3 and CMT2 in green plants (*Viridiplantae*). (B) The protein sequence identity between CMT2 and CMT3 (left) and the residue contacting +2 nucleotide of CHG and corresponding residue in CMT2 (right) in flowering plants. (C) *In vitro* DNA methyltransferase assay of CMT2 and CMT2^V1200R^ on CHH, CHG or hemi-methylated CHG (hmCHG) DNA substrates. (D) Structure of Maize ZMET2-DNA complex (PDB 7UBU) showing the interaction between ZMET2 Y526 and R804 and the guanine at the +2 position (G_+2_) from the target cytosine in a CHG context. The ZMET2 R804-corresponding residue in CMT3, R745, is shown in parenthesis. (E) Structural model of *Arabidopsis* CMT2 showing the position of G_+2_ and its surrounding residues, V1200 and H924. (F) Phenotype images of two independent transgenic CMT2 and CMT2^V1200R^ plants in *cmt2cmt3* (*cc*) mutant background. (G) Immunoblots showing CMT2 and CMT2^V1200R^ protein levels from the plants in (F). Actin serves as a loading control. (H) McrBC-qPCR assay determining DNA methylation levels of two CMT2-targeted TEs, *Copia89* and *Copia18A*. (I) Genome browser view of CHG and CHH methylation levels of *Copia89* from bisulfite-sequencing data. (J) Metaplots showing the average CHG methylation over all TEs in *Arabidopsis*. (K) Venn diagram showing the overlapping of DMRs identified by comparing indicated genotypes with Col-0. See also Supplementary Figure 1-5 and Supplementary Dataset 1.

CMT2 is paralogous to CMT3 and shares high protein sequence similarity, especially in basal angiosperms (Figure 1B). Interestingly, they demonstrate distinct substrate preference for DNA methylation. While CMT3 preferentially methylates hemi-methylated CHG ^20,21^, CMT2 exhibited a considerably higher methylation activity on CHH over CHG DNA substrate (Figure 1C and Supplementary Figure 3A). In addition, CMT2 showed a modest (∼2-fold) substrate preference for hemi-methylated over unmethylated CHG DNA (Figure 1C and Supplementary Figure 3A), suggesting a residual maintenance methyltransferase activity of CMT2 for CHG sites, in line with previous reports of redundancy ^14,17^. To further understand the CMT2 evolution, we performed a similar phylogenetic analysis and found that CMT2 from basal angiosperms *Amborella trichopoda* contains the same arginine in the equivalent to CMT3 R745, a key residue for CHG substrate recognition (Figure 1B). However, this arginine has diversified into different non-positively charged residues in higher angiosperm species such as cysteine in *Setaria italica*, phenylalanine in *Glycine max*, and valine in *Arabidopsis thaliana* (Figure 1B and Supplementary Figure 1B). The conservation of this CHG-recognizing arginine residue in basal angiosperms further supports the notion that CMT2 originates from CMT3 duplication, likely along with the whole genome duplication event epsilon (ε) during angiosperms evolution (Figure 1B) ^25^.

### CMT2 V1200R induces a gain in CHG methylation *in vitro* and *in vivo*

To investigate whether this change of CHG-recognizing arginine causes the loss of CHG specificity in CMT2, we first generated a structural model of the CMT2 in complex with hmCHG (Supplementary Figure 3B-F) based on the ZMET2 crystal structure. We found that the hydrogen-bonding interaction for base-specific recognition of the +2 guanine (G_+2_) in the ZMET2-hmCHG complex was no longer preserved in CMT2, due to replacement of ZMET2 Y526 and R804 by the corresponding H924 and V1200 of CMT2 (Figure 1D,E), explaining the loss of the CHG specificity of CMT2. We then substituted the corresponding V1200 in *Arabidopsis thaliana* CMT2 with arginine (CMT2^V1200R^). Our *in vitro* methyltransferase assay showed that the CMT2^V1200R^ mutation has activities about 16-fold higher on hmCHG and 8-fold higher on CHG substrates compared to the wild-type CMT2 (Figure 1C). To understand the effect of this mutation *in vivo*, we expressed the genomic fragments of CMT2 and CMT2^V1200R^ fused with a FLAG tag in *cmt2 cmt3* (*cc*) and *drm1 drm2 cmt2 cmt3* (*ddcc*) mutants (Figure 1F,G, and Supplementary Figure 4A,B). We first investigated the DNA methylation levels of two TEs co-targeted by CMT2 and CMT3, *Copia89* and *Copia18A*, by McrBC-qPCR assay, and found the introduction of V1200R led to higher methylation than wild-type CMT2 (Figure 1H and Supplementary Figure 4C). We next performed bisulfite-sequencing to investigate the effect of the V1200R mutation on whole-genomic methylation levels (Supplementary Table 1). While complementation by CMT2 led to a partial rescue of CHG methylation, introduction of V1200R led to a gain of global CHG methylation level with or without the *de novo* methyltransferase DRM2 (Figure 1I,J, and Supplementary Figure 4D-G).

To test whether the V1200R mutation could alter CMT2 targeting, we compared the differentially methylated regions (DMRs) and examined their genomic locations. We found that most TEs targeted by CMT2^V1200R^ for CHG methylation were original CMT2 targets for CHH methylation (Figure 1K). To further investigate the chromatin features of these regions that gained CHG methylation by CMT2^V1200R^, we explored their local chromatin environments and locations in the genome. We found that like CMT2, CMT2^V1200R^ methylated regions contain lower chromatin accessibility and enriched with H3K9me2 and H3K27me3 (Supplementary Figure 5A-E). We also found that these regions with increased CHG methylation by CMT2^V1200R^ corresponded to chromatin states annotated for intergenic and centromeric regions (Supplementary Figure 5F). Taken together, these data suggested that CMT2^V1200R^ mutation renders gain of CHG methylation function with minimal effect on chromatin targeting.

### CMT2^V1200R^-mediated CHG methylation represses TEs

To understand the transcriptional consequences from the gain of CHG methylation by CMT2^V1200R^, we performed RNA-seq in CMT2/*cc* and V1200R/*cc* with Col-0 and *cc* as controls (Supplementary Table 2). We noted that a subset of TEs (n = 42) upregulated in *cc* due to hypomethylation were re-silenced by the introduction of CMT2^V1200R^ but not CMT2 (Figure 2A). These TEs were enriched for *LINE*/*L1* and *LTR*/*Copia* type retrotransposons (Supplementary Figure 6A). We further determined the DNA methylation level in TEs that were specifically re-silenced by V1200R/*cc* and found that the CHG methylation levels were comparable to Col-0 in V1200R/*cc*, while CMT2 only partially re-methylated these TEs in *cc* (Figure 2B), suggesting that the low expression in V1200R/*cc* was due to gain of CHG methylation. We next examined the expression of two TEs, *Copia11* and *Copia47*, and found that both TEs were re-silenced by CMT2^V1200R^, but not CMT2 (Figure 2C).

**Figure 2.**
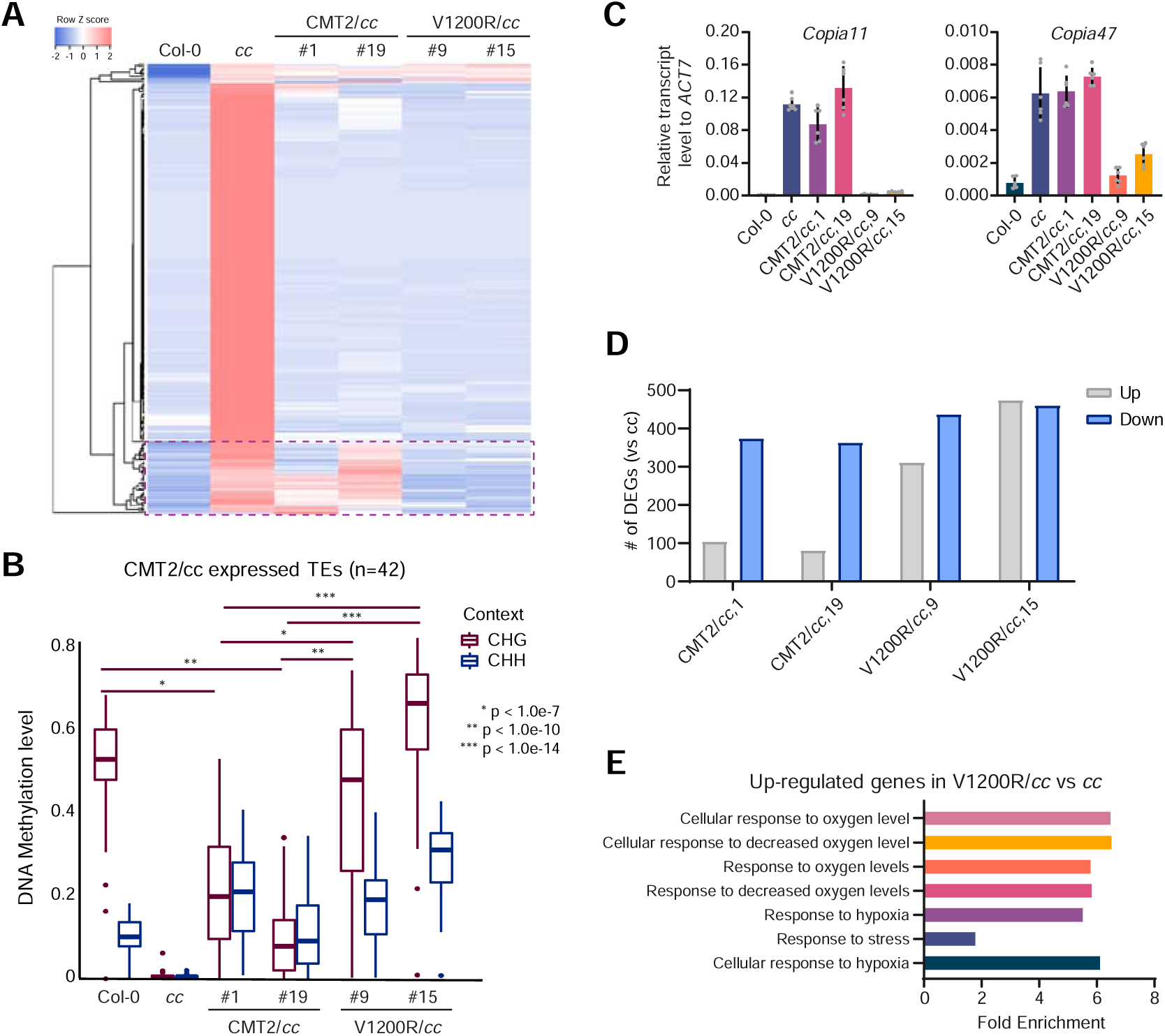
CMT2^V1200R^ represses both CMT2- and CMT3-targeted TEs. (A) Heat maps showing expression levels of up-regulated TEs in two independent CMT2/*cc* and V1200R/*cc* transgenic lines. *cc*: *cmt2cmt3*. The dashed box indicates TEs re-silenced by CMT2^V1200R^. (B) CHG and CHH DNA methylation levels of TEs remaining expressed in CMT2/*cc*. (C) Relative transcript levels of *Copia11* and *Copia47* measured by RT-qPCR in CMT2/*cc* and V1200R/*cc* transgenic plants. The transcript level was normalized to *ACT7*. (D) Number of differentially expressed genes (DEGs) in CMT2/*cc* and V1200R/*cc*. (E) Gene Ontology terms of upregulated genes in V1200R/*cc*. See also Supplementary Figure 6.

Similarly, we found more differentially expressed genes (DEGs) in the two V1200R/*cc* lines compared with CMT2/*cc* (Figure 2D). Among the common DEGs of two V1200R/*cc* lines, many of the upregulated genes (n=269) have low expression levels in Col-0, *cc*, and CMT2/*cc* (Supplementary Figure 6B). Conversely, lowly expressed genes (n=85) in V1200R/*cc* have higher expression in Col-0, *cc*, and CMT2/*cc* (Supplementary Figure 6B). Genes upregulated in only V1200R/*cc* are enriched for gene ontology terms involved in response to stress and decreased oxygen levels (Figure 2E). Elevated expression of stress-related genes in V1200R lines may help explain the developmentally stunted, smaller plant phenotype (Figure 1F).

### CMT2 contains a long N-terminus important for nuclear localization

Despite sharing similar protein domains, CMT2 and CMT3 differ in their protein lengths with CMT2 possessing a longer N-terminus compared to CMT3 in *Arabidopsis* (i.e. ∼550 aa in CMT2 and ∼90 aa in CMT3) (Figure 3A). Similarly, *Amborella* CMT2 (AmtriCMT2) contains a long N-terminus (Figure 3A) despite sharing similar domain structures with AmtriCMT3 (Figures 3A and 1B). Further analysis revealed a longer N-terminus of CMT2 compared to CMT3 in representative angiosperm species that contain both proteins, suggesting a conserved function of this long N-terminus (Supplementary Figure 7A). Interestingly, we found that AmtriCMT2 N-terminus contains eight tandem repeats, each encoding conserved ‘RRSPR’ sequences (Figure 3B and Supplementary Figure 7B). This ‘RRSPR’ motif is conserved in CMT2 of other angiosperm species, varying from one to nine copies containing RRSxR (x represents any amino acid) (Supplementary Figure 7C). To investigate how the long N-terminus have emerged, we compared the DNA sequences of AmtriCMT2 and AmtriCMT3 and found that the AmtriCMT2 N-terminus sequence bared similarity to AmtriCMT3 promoter sequences (Supplementary Figure 8A), suggesting that the long N-terminus of AmtriCMT2 may arise from exonization of AmtriCMT3 promoter during a duplication event (Supplementary Figure 8B).

**Figure 3.**
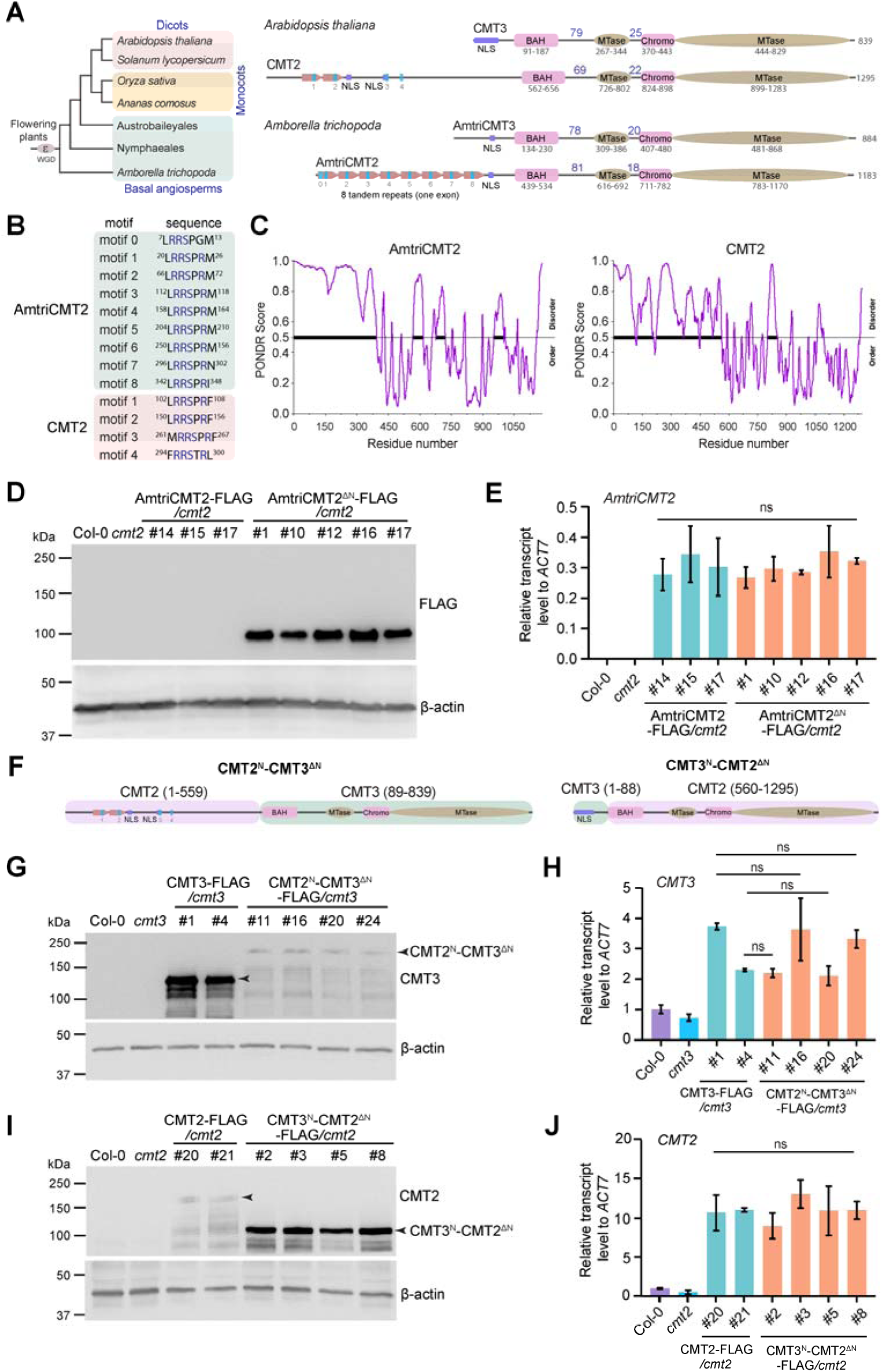
The long CMT2 N-terminus is disordered and controls protein level. (A) Diagrams showing the domain structure of CMT2 and CMT3 from *Arabidopsis thaliana* (dicots) and *Amborella trichopoda* (basal angiosperms). The blue bars indicate tandem repeats. NLS, nuclear localization sequence. Numbers above lines depict the number of amino acids between domains. (B) Alignment of the conserved RRS motifs in CMT2 N-terminus of *Amborella* and *Arabidopsis*. (C) Graph of intrinsic disorder scored by PONDR for CMT2 of *Amborella* and *Arabidopsis*. Y-axis indicates PONDR VSL2 scores and x-axis indicates the residue positions. (D) Immunoblots for protein levels of full length *Amborella* CMT2 (AmtriCMT2) and N-terminus-truncated *Amborella* CMT2 (AmtriCMT2^ΔN^) expressing in *Arabidopsis cmt2* mutant background. Actin serves as a loading control. Numbers represent the independent transgenic lines. (E) RT-qPCR showing relativ*e AmtriCMT2* transcript levels in AmtriCMT2 and AmtriCMT2^ΔN^ transgenic lines normalized to *ACT7*. (F) Diagrams showing the swapping of N-terminus between CMT2 and CMT3. Residue numbers were depicted above the sections. (G) Immunoblots for protein levels of wild type (CMT3) and N-terminus swapped CMT3 (CMT2^N^-CMT3^ΔN^) transgenic lines in *cmt3* mutant. (H) RT-qPCR showing relativ*e CMT3* transcript levels in CMT3 and CMT2^N^-CMT3^ΔN^ transgenic plants. Transcript levels were first normalized to *ACT7* and then to *CMT3* level in Col-0. (I) Immunoblots for protein levels of wild type (CMT2) and N-terminus swapped CMT2 (CMT3^N^-CMT2^ΔN^) transgenic lines in *Arabidopsis cmt2* mutant. (J) RT-qPCR showing relativ*e CMT2* transcript levels of CMT2 and CMT3^N^-CMT2^ΔN^ transgenic lines. Transcript levels were first normalized to *ACT7* and then to *CMT2* level in Col-0. See also Supplementary Figure 7-9.

Next, we predicted AmtriCMT2 N-terminus structure and found it to be disordered using PONDR (Figure 3C). The N-terminus of both CMT3 and CMT2 is also predicted to contain nuclear localization sequences (NLS) (Figure 3A, Supplementary Figure 9A). Upon deletion of the N-terminus, we observed a cytosolic localization of CMT3 in both *N. benthamiana* and *Arabidopsis* (Supplementary Figure 9B, C). Similarly, CMT2 N-terminus deletion (CMT2^ΔN^-GFP) resulted in cytoplasmic localization in both *N. benthamiana* and *Arabidopsis*, whereas the N-terminus only (CMT2^N^-GFP) showed nuclear localization (Supplementary Figure 9B, C). Our nuclear fractionation experiment further confirmed the importance of N-terminus for CMT2 nuclear localization (Supplementary Figure 9D).

### The N-terminus of CMT2 regulates protein stability

To investigate the N-terminus function, we generated transgenic *Arabidopsis* plants expressing the full-length AmtriCMT2 as well as an N-terminally truncated AmtriCMT2 without the tandem repeats while retaining the NLS (AmtriCMT2^ΔN^, aa 373-1183). Surprisingly, we noted a much higher AmtriCMT2^ΔN^ protein abundance than that of AmtriCMT2, despite similar transcript levels (Figure 3D,E). We next asked whether this observed N-terminus function of AmtriCMT2 was shared by *Arabidopsis* CMT2. Instead of truncation, we swapped the N-terminus of *Arabidopsis* CMT2 and CMT3 with consideration for the essential role of the NLS (Figure 3F and Supplementary Figure 9). We transformed CMT3 and CMT2^N^-CMT3^ΔN^ into *cmt3* mutant and found that fusion of CMT2 N-terminus resulted in much lower CMT3 protein level despite having similar transcript levels (Figure 3G,H). Consistently, CMT2 protein level was much higher when the long N-terminus was replaced by the shorter CMT3 N-terminus (Figure 3I,J). These results suggest that the long N-terminus regulates CMT2 protein level.

We next performed cycloheximide (CHX, a protein synthesis inhibitor) experiment to directly test the CMT2 protein stability. We found that CMT2^N^-CMT3^ΔN^ protein level decreased quickly and was halved after 4 hours of CHX treatment, whereas CMT3 remained relatively stable and decreased at a much slower rate (Figure 4A,B). Inversely, CMT2 protein level was halved after 6 hours after CHX treatment, while the CMT3^N^-CMT2^ΔN^ protein remained stable (Figure 4B,C). Interestingly, the N-terminus alone (CMT2^N^-GFP) was sufficient to affect protein stability in a similar decrease pattern as CMT2^N^-CMT3^ΔN^ (Figure 4D). Altogether, these data showed that the N-terminus of CMT2 regulates its protein stability.

**Figure 4.**
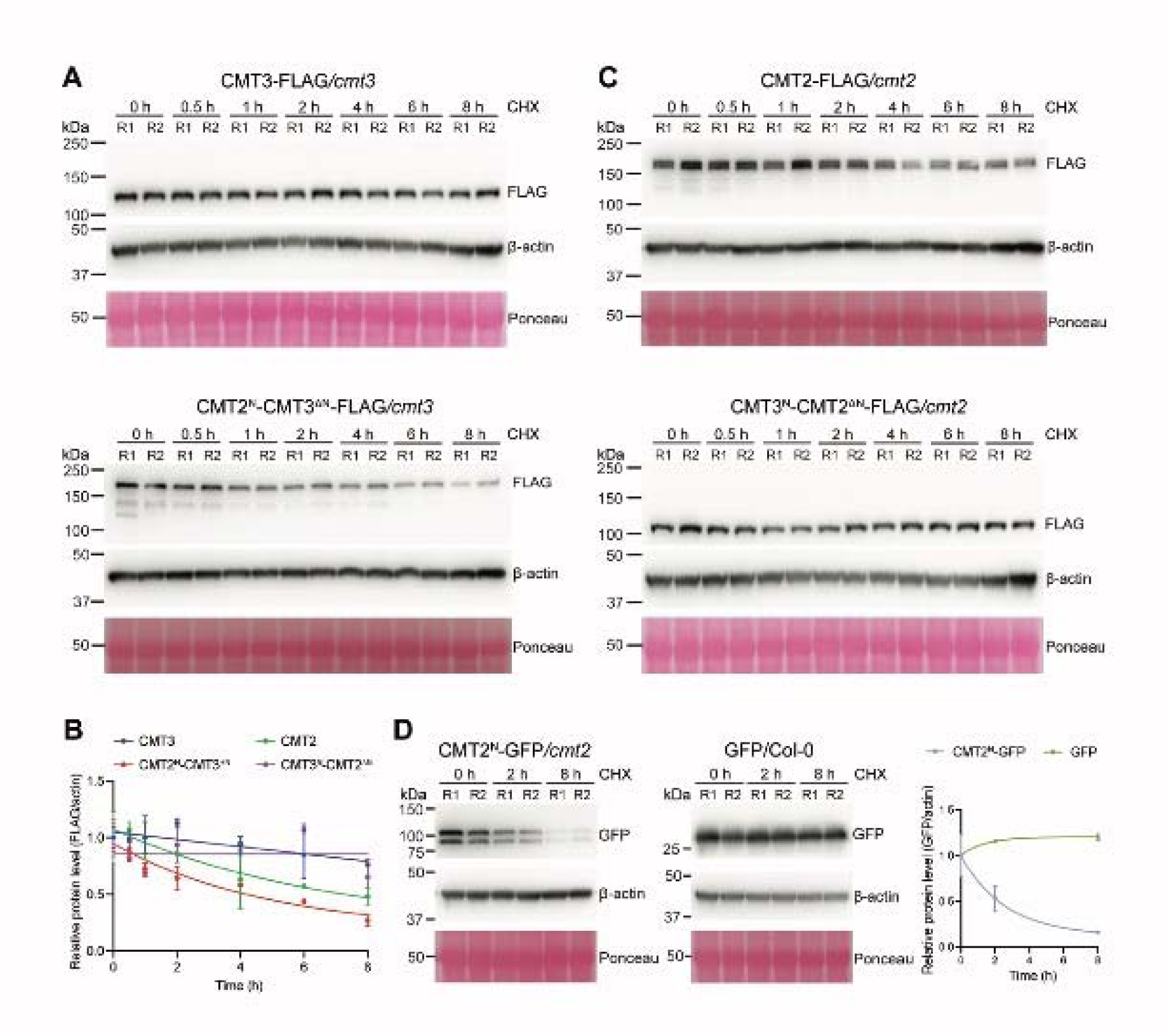
CMT2 N-terminus regulates its protein stability. (A) Immunoblots showing the levels of CMT3 (top) and CMT2^N^-CMT3^ΔN^ (bottom) after cycloheximide (CHX, 500 μM) treatment for the indicated time. 7-day-old seedlings were used. Actin served as a control. R1 and R2 represent two biological replicates. (B) Quantification of protein levels normalized to actin. (C) Immunoblots showing the levels of CMT2 and CMT3^N^-CMT2^ΔN^ after treatment with 500 μM CHX for the indicated time. (D) Immunoblots showing levels of CMT2^N^-GFP (left) and GFP only (middle) after treatment with 500 μM CHX. Quantification of protein levels by GFP signals normalized to actin (right).

### N terminus mediates heat-induced CMT2 degradation

We showed previously that CMT2 was degraded upon heat stress and CMT3 remained stable ^17^. To determine whether CMT2 N-terminus mediates the heat-induced protein instability, we exposed our transgenic swapped lines to 37 °C treatment and found that the fusion of CMT2 N-terminus to CMT3 (CMT2^N^-CMT3^ΔN^) demonstrated a much higher sensitivity to heat, where protein level was halved after 6 hours of heat treatment compared to the stable protein level in wildtype CMT3 (Figure 5A,B). Conversely, the fusion of N-terminus of CMT3 to CMT2 (CMT3^N^-CMT2^ΔN^) greatly increased protein stability under heat stress compared to the fast degradation of wildtype CMT2 (Figure 5B,C). Consistently, CMT2^N^-GFP protein abundance decreased and was undetectable after 24 hours of heat treatment (Figure 5D). Furthermore, we observed reduced nuclear localization of CMT2^N^-GFP upon 4 hours of heat treatment, similar as the full-length CMT2 (Figure 5E). Together, these results demonstrated that N-terminus mediates heat-induced CMT2 degradation.

**Figure 5.**
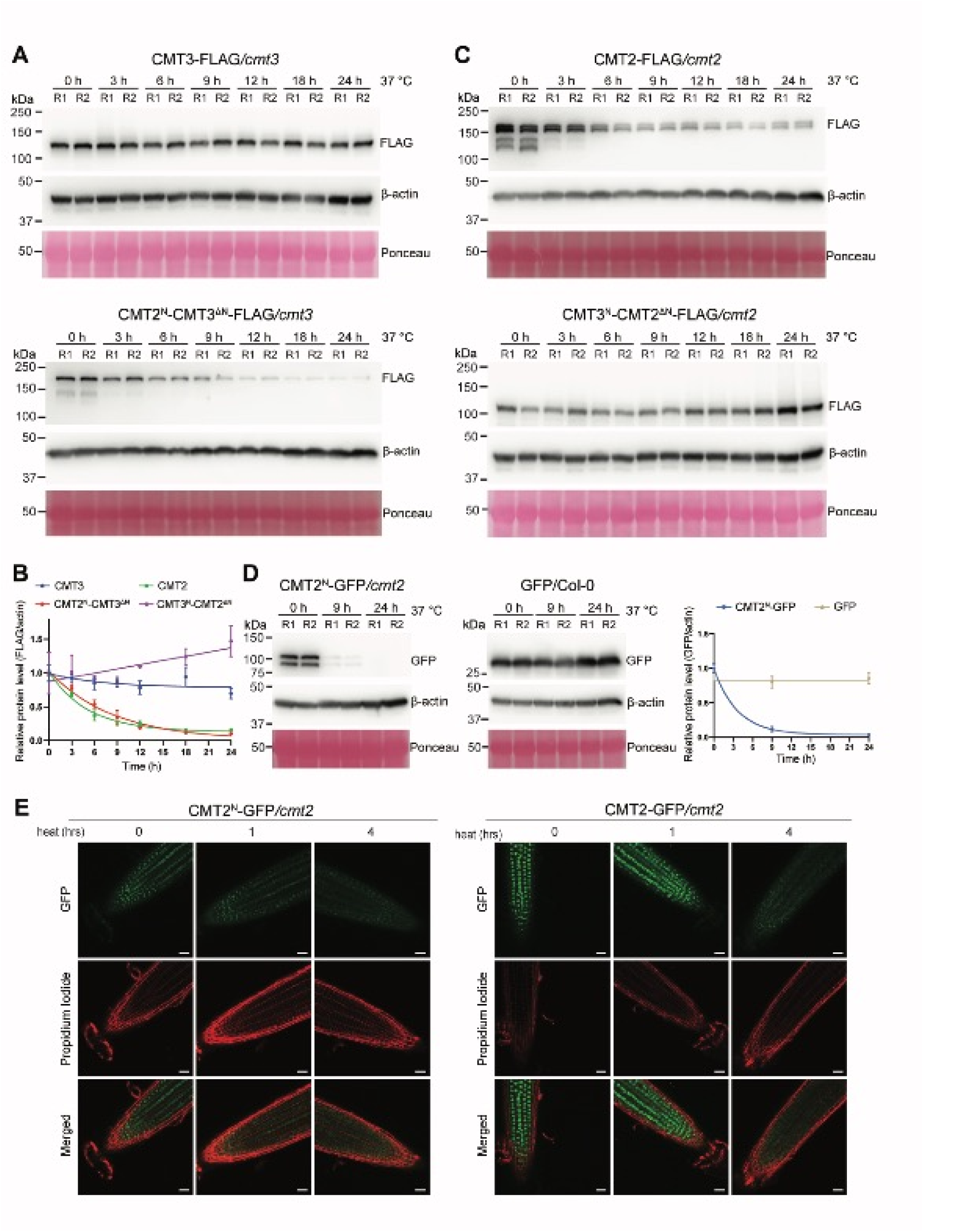
The long N-terminus mediates heat induced CMT2 protein degradation. (A) Immunoblots detecting the levels of CMT3 and CMT2^N^-CMT3^ΔN^ after 37°C heat treatment for the indicated time. 10-day-old seedlings were used. Actin served as a control. R1 and R2 represent two biological replicates. (B) Quantification of protein levels normalized to actin. (C) Immunoblots detecting the levels of CMT2 and CMT3^N^-CMT2^ΔN^ after 37°C heat treatment for the indicated time. (D) Immunoblots showing levels of CMT2^N^-GFP (left) and GFP only (middle) after 37 °C heat treatment. Quantification of protein levels by GFP signals normalized to actin (right). (E) Confocal images showing the localization and intensity of CMT2^N^-GFP and CMT2-GFP in *Arabidopsis* root tip upon 37°C heat treatment for the indicated time (scale bar: 25 μm).

### Natural variation of CMT2 demonstrated tolerance to environmental stress

Many *Arabidopsis* natural accessions with CHH methylation variations have been linked to CMT2 ^18,26-29^. Considering the disordered nature of the CMT2 N-terminus, we wonder whether the N-terminus is more plastic to harbor natural mutations. We examined the CMT2 variations in the 1001 genome project ^30^ and found that the variations were indeed mostly located at the N-terminus (Supplementary Figure 10A and Supplementary Dataset 2). Our *dN*/*dS* analysis on the CMT2 coding sequences among close relatives of *A. thaliana* in the Brassicaceae family also showed that the N-terminal codons appeared to be evolutionarily neutral (Figure 6A). Upon re-analysis of BS-seq data from 48 accessions with CMT2 nonsense or frameshift mutations from the 1001 epigenome project ^18^, we found that global CHH methylation level varied with the mutation position in CMT2 and the geographical location of these accessions (Figure 6B and Supplementary Figure 10B). Stop-gain mutations in the first two exons of CMT2 are relatively common in nature and appear not to affect genome-wide methylation but may rather increase CHH methylation variability (Figure 6B,C). Meanwhile, mutations in the C-terminal methyltransferase domain are much less commonly detected and have generally lower methylation relative to the N-terminal mutations (Figure 6C).

**Figure 6.**
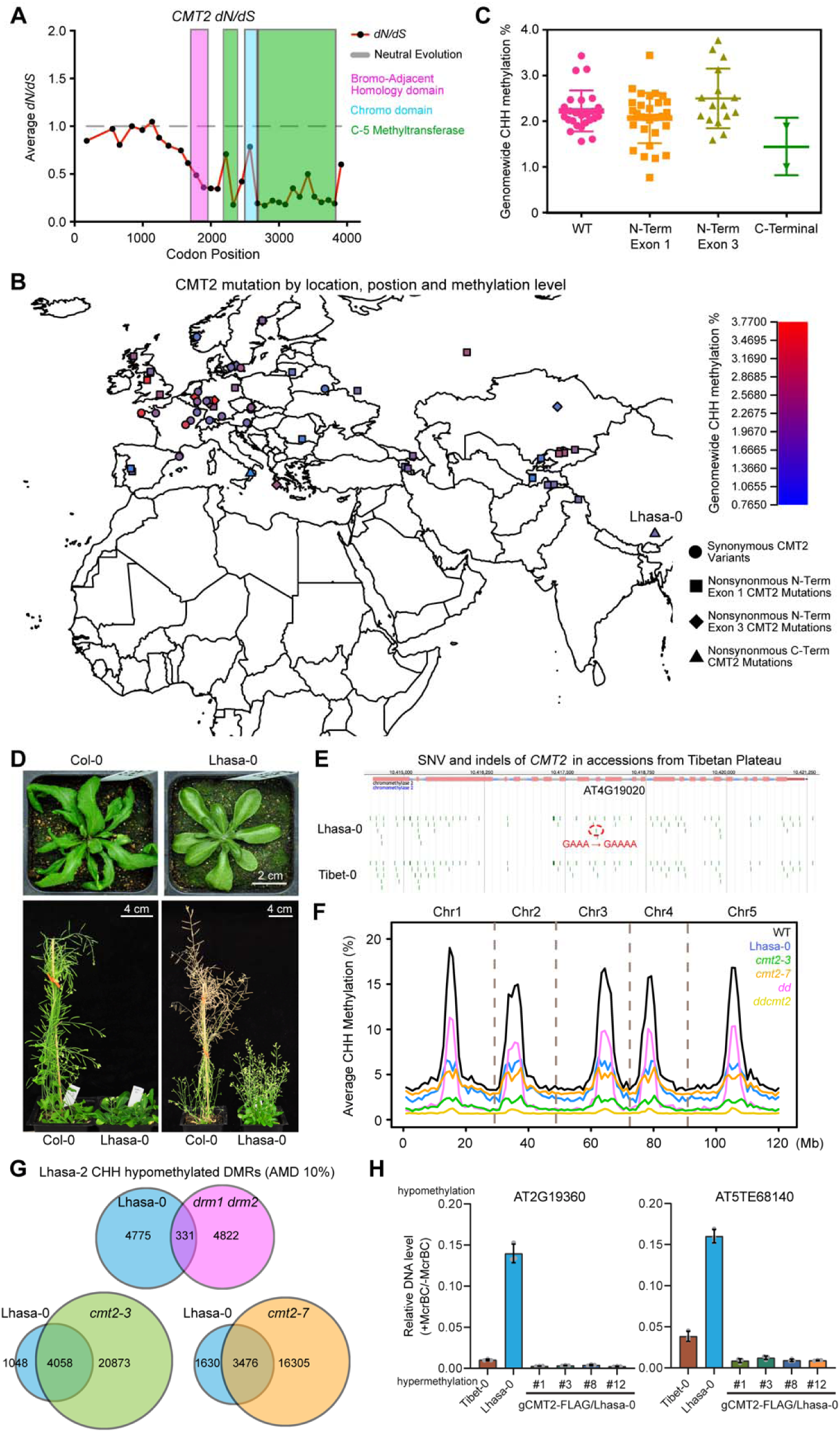
Natural variations of CMT2 in *Arabidopsis thaliana*. (A) *dN*/*dS* plot of *CMT2* gene in *Brassicaceae*. (B) Genome-wide CHH methylation levels of accessions with different mutations in *CMT2* gene. (C) Dot plot of CHH methylation in WT (n=25), N-terminal Exon 1 (n=31), N-terminal Exon 2 (n=16), and C-terminal (n=2) CMT2 mutations. (D) Phenotype comparisons between *Arabidopsis* ecotypes Col-0 and Lhasa-0. (E) Genome browser view of natural accessions from the Tibetan Plateau, Lhasa-0 and Tibet-0, showing single nucleotide variant (SNV) with an insertion in the CMT2 exon. (F) Average CHH methylation levels across *Arabidopsis thaliana* chromosomes of Lhasa-0 ecotype in comparison with wildtype Col-0 and mutants of CHH DNA methyltransferases. (G) Overlapping of hypomethylated DMRs between Lhasa-0 and mutants of CHH DNA methyltransferases. (H) McrBC-qPCR assay showing DNA methylation complementation at two representive loci. The transgenic lines with CMT2 genomic sequence (gCMT2) can rescue lost CHH methylation in Lhasa-0. Tibet-0 was used as a control. See also Supplementary Figure 10-11 and Supplementary Dataset 2.

To gain additional insights into CMT2 natural variation and CHH methylation, we analyzed *Arabidopsis thaliana* accessions from various locations. Interestingly, we found a natural accession, Lhasa-0, contains a CMT2 mutation in the methyltransferase domain. Lhasa-0 was isolated from the Tibetan plateau (altitude 4200m), closely located to Tibet-0 accession (Figure 6B and Supplementary Figure 11A)^31^. Lhasa-0 plants have a broader, more circular leaf structure, dwarf, much more branching, and late flowering compared to Col-0 (Figure 6D and Supplementary Figure 11B,C). We sequenced the genome of Lhasa-0 and found it contains one bp insertion at the 10^th^ exon of CMT2, resulting in a frameshift mutation in methyltransferase domain and premature stop codon (Figure 6E, Supplementary Figures 10B and 11D,E). Next, we performed BS-seq and found global CHH methylation reduction in Lhasa-0, to a similar degree of *cmt2* mutants (Figure 6F). We next called the DMRs using the hcDMR calling program ^32^ and found many hypomethylated CHH DMRs (n=5106) in Lhasa-0 (Figure 6G). These hypo-DMRs overlapped significantly with *cmt2* but not *drm1drm2* (Figure 6G), suggesting a link between CMT2 and CHH methylation in Lhasa-0. Furthermore, we found that methylation at the two hypomethylated genes could be rescued by introduction of the genomic fragment of functional *CMT2* transgene into Lhasa-0 (Figure 6H). These data suggest that CMT2 is responsible for the CHH hypomethylation phenotype of Lhasa-0.

To determine if mutations in CMT2 could confer any phenotypic advantages or consequences to wild accessions, we performed stress tolerance assays with Col-0, Lhasa-0, and *cmt2* mutants of Col-0. Lhasa-0 showed stronger tolerance to UV-B stress, while *cmt2-7* also survived in UV-B stress at a slightly higher rate than Col-0 (Supplementary Figure 11F,G). In addition, Lhasa-0, and two *cmt2* mutants show resistance to heat stress (Supplementary Figure 11H,I), in agreement with a previous report ^29^. Collectively, these data suggest that CMT2 may be involved in plant survival under environmental stresses.

## Discussion

In this study, we revealed the mechanisms diversifying two members of the CMT family, CMT2 and CMT3, for diverse heterochromatic non-CG methylation in plants. According to structural model and biochemical assays, CMT2 and CMT3 share a similar enzymatic mechanism for non-CG methylation, ensuring their overlapped functionalities in non-CG methylation (Figure 7A). However, an arginine residue that mediates recognition of +2 guanine of CHG via a hydrogen-bonding interaction is diversified in angiosperm CMT2, providing an explanation of why CMT3 and CMT2 show distinct preference for CHG and CHH methylation, respectively ^21^. Consistently, a single V1200R mutation of CMT2 gains maintenance activity for both CHG and CHH methylation in *Arabidopsis* (Figure 7A). During evolution, CMT2 may have evolved from a whole genome duplication event before the emergence of angiosperms about 181 million years ago, as evidenced by the retention of the key arginine CMT3 residue in the CMT2 of basal angiosperm species *Amborella trichopoda* (Figure 7B). However, CMT2 is different from CMT3 since its birth, where its N-terminus becomes longer, due to a tandem repeat insertion and exonization (Figure 7B). The long N-terminus is disordered, undergoes neutral selection during evolution, and makes the protein relatively unstable. Nevertheless, the N-terminus contains several conserved tandem RRS motifs, though the number and position vary in flowering plants. The arginine variation together with the long N-terminus drives the evolution of CMT2 for CHH methylation preference.

**Figure 7.**
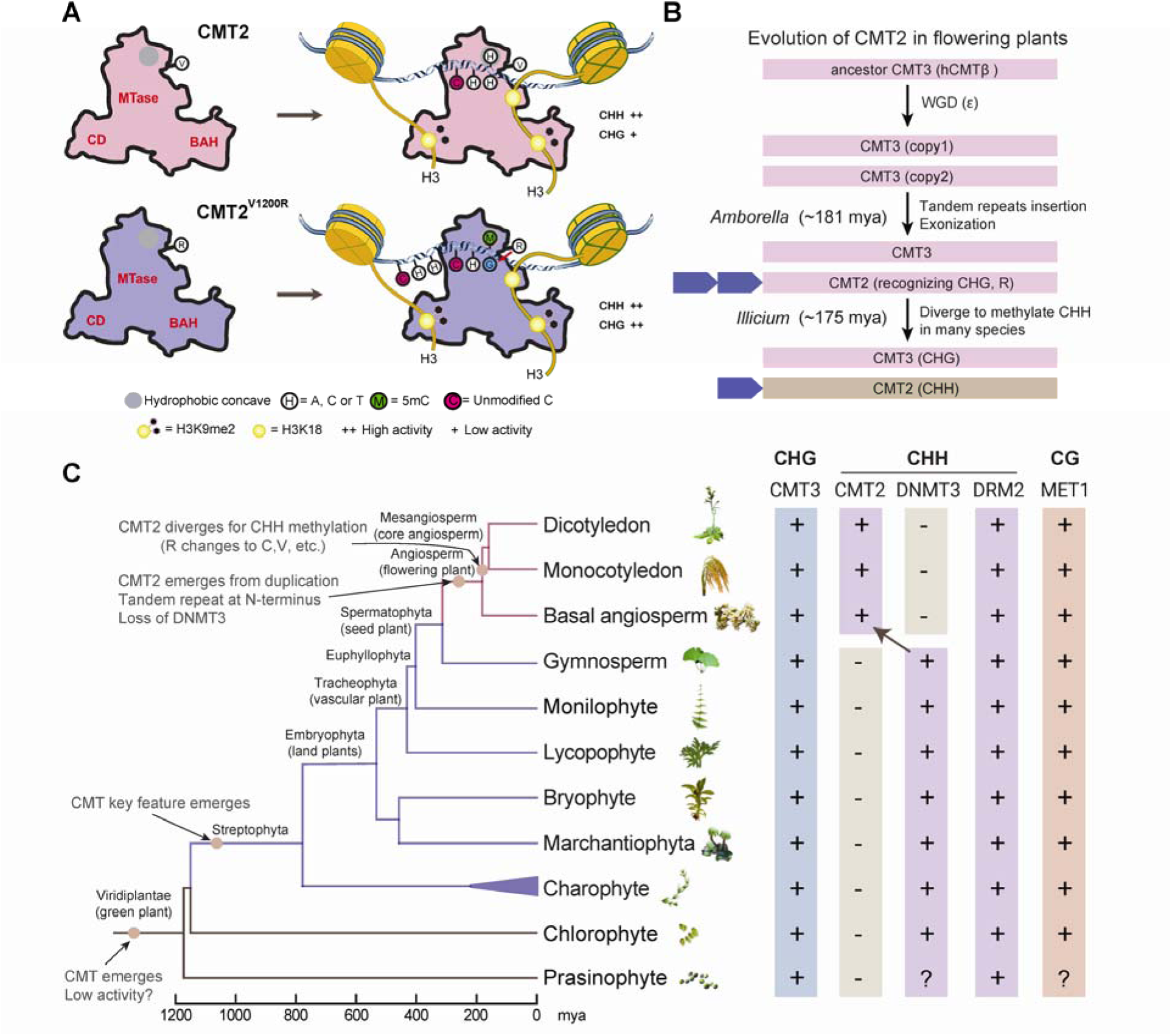
A working model for the evolution of chromomethylases in plants. (A) A mechanistic model for the enzymatic regulation and specificity of CMT2 and CMT2^V1200R^ in the chromatin environment. (B) Proposed model of CMT2 emergence and evolution in the flowering plants. (C) Phylogenetic tree depicting predicted emergence and divergence of chromomethylases in plant (left) and DNA methyltransferases found in the representative plant species (right). See also Supplementary Figure 12.

It is worth noting that recent studies have indicated that DNMT3s, which are homologous to mammalian DNMT3A and DNMT3B, also play a role in CHH methylation maintenance in heterochromatic regions in some plant species such as the moss *Physcomitrella patens* ^23,33^. Intriguingly, DNMT3s are widely present in the plant kingdom, but have become extinct in angiosperms (Figure 7C). In this context, the evolutionary emergence of CMT2 presumably provides a similar, possibly even more robust, mechanism in replacing the loss of DNMT3-mediated CHH methylation and maintaining the non-CG methylation at heterochromatin (Figure 7C). The functional context behind the evolutionary transition between CMT2 and DNMT3s awaits further investigation.

CMT2 appears to be more plastic and could offer a great target for evolution, where it contains much more variations in nature or is even lost in certain angiosperm species. In maize, CMT2 is lost and the CMT3 (ZMET2) has both CHG and CHH methylation activity, yet whether this could be a reason for the high TE abundance and activity remains to be investigated ^9,34^. It was previously reported that TEs in *A. thaliana* natural accessions varied greatly, corresponding to differences in gene expression and DNA methylation in these accessions, which could provide a source for genetic diversity in the population ^35^. We found an abundance of natural variations in CMT2, especially in the N-terminus. We also identified a new *Arabidopsis thaliana* accession Lhasa-0 carrying a CMT2 mutation leading to global DNA hypomethylation. As Lhasa-0 is isolated from Tibetan Plateau, a high-altitude environment with harsh conditions such as strong sunshine and UV-B radiation, the mutation in CMT2 could possibly confer some tolerance to environmental stresses ^31,36^. Thus, over evolutionary time, mutations in CMT2 could potentially alleviate repression on TEs which could then increase genetic diversity. Consistent with this idea, we found another species, *Arabis alpina*, a close relative of *A. thaliana* from high altitude alpines, also contains a CMT2 mutation with the loss of the essential BAH domain (Supplementary Figure 12). TEs, especially retrotransposons, have also been expanded, accompanied with altered DNA methylation in *Arabis alpina* ^37^. The transcriptional activity of TEs also seems to increase along with an elevational gradient in *Arabidopsis arenosa* ^38^. Previously, we found that UV-B also triggered DNA hypomethylation via the interaction between UVR8 and DRM2 ^39^. Whether CMT2 is also involved in UV-B suppressed DNA methylation requires further investigation.

CMT2 stability is sensitive to heat stress, mainly conferred by its long and disordered N-terminus. Though the N-terminus varies in different angiosperm species, many contain the conserved RRS motifs, indicating that these motifs may have yet undiscovered functions. Degradation of CMT2 under high temperature could impede its function for CHH methylation maintenance at heterochromatic TEs, thereby alleviating some TEs that were normally suppressed under normal conditions. Over the course of evolution, some de-repressed TEs could potentially jump in the genome and generate various genetic mutations for natural selection, which could be especially beneficial for species survival under stress. Indeed, the variation of TE transposition in *A. thaliana* wild accessions is related to climate change, where the increase in annual temperature correlates with an increase of *AtCopia87* (*ONSEN*) copy number ^40^. As another support, natural CMT2 variation has been associated with genome-wide methylation changes and temperature seasonality ^29^. During the earth history and in a large period, the average global temperature is much higher (up to more than 10°C) than that of today ^41^. Thus, we can infer that during the hothouse era, the instability of CMT2 together with the TEs may have helped shape the plant evolution on earth.

## Materials and Methods

### Plant materials, plasmid construction, and transformation

*Arabidopsis thaliana* ecotypes Columbia-0 (Col-0), Tibet-0 and Lhasa-0 (gifted from Professor Yang Zhong) were used in this study. The Col-0 ecotype was used as the background for all mutant and transgenic plants, except for experiments in the Lhasa-0 ecotype. Mutant lines used in this study were *cmt3* (*cmt3-11*, SALK_148381), *cmt2 (cmt2-3* SALK_012874 and *cmt2-7,* WiscDsLox7E02), *cc* (*cmt2-7 cmt3-11, cmt2-7* is WiscDsLox7E02), and *ddcc* (*drm1-2 drm2-2 cmt3-11 cmt2-7*). All seedlings and plants used in this study were grown in long-day conditions (16/8-hr light/dark cycle) at 22 °C in a Conviron chamber. *Nicotina benthamiana* plants were grown directly on soil and under long-day conditions.

Genomic fragments of *Arabidopsis* CMT2 or CMT3 were used to construct plasmids for transgenic lines. Using genomic DNA extracted by the CTAB method as the template, genomic fragments including the native promoter (CMT2,1.4 kb; CMT3, 1.6 kb), 5’-UTR, exons, and introns were amplified by PCR, and then ligated to linearized vector by in-fusion cloning method (Vazyme, C115). For FLAG-tagged lines, the vector pCAMBIA-FLAG-FAST containing a 3x FLAG tag at the C-terminus of the genomic fragment and a selection marker pOLE1:OLE1-RFP, which is specifically expressed in dry seeds, was used ^42^. For GFP-tagged lines, the vector pCAMBIA1302 containing an *mgfp5* marker gene was used. For the expression of CMT2 and CMT3 of other plant species e.g. *Amborella trichopoda* (from Botany Garden and Greenhouses of UW-Madison, *Chara brauii* (from Kobe University), and *Marchantia polymorpha* (collected from Madison, WI), the protein-coding sequence was amplified from cDNA and fused after the UBQ10 promoter and ligated to linearized pCAMBIA-FLAG-FAST vector.

Transgenic lines used in this study include *gCMT3-FLAG, gCMT3-GFP,* domain truncation and swapping generated in *cmt3* and *cc* backgrounds; *gCMT2-FLAG*, *gCMT2^V1200R^-FLAG, gCMT2-GFP*, domain truncation and swapping generated in *cc*, *cmt2, cmt3,* and *ddcc* backgrounds; CMT2 and CMT3 of other species were generated in respective *Arabidopsis cmt2* and *cmt3* backgrounds. Transgenic plants were generated via floral-dip method and screened with a hand-held fluorescent lamp (Luyor-3415RG) for OLE1-RFP expression in seeds or by hygromycin selection. For CMT2 complementation in Lhasa-0 background, genomic CMT2 fragment from Col-0 were transformed into Lhasa-0 plants using similar methods as above.

### Immunoblotting

Frozen plant samples were ground to powder with a tissue lyser (Scientzbio). Proteins were extracted by boiling sample in 5% SDS at 95 °C for 10 mins. Proteins were separated by SDS-PAGE and transferred to nitrocellulose membranes. Membranes were blocked with blocking buffer (5% non-fat milk in TBST) for 1 hour at room temperature, and then incubated with primary antibodies (diluted in 3% BSA in TBST) overnight at 4 °C. For unconjugated primary antibodies, membranes were incubated with HRP-conjugated secondary antibodies in blocking buffer for 1 hour at room temperature. Membranes were washed with TBST for 5 mins, three times, before imaging. ECL substrate was mixed and added to the membrane prior to imaging, and chemiluminescence images were taken with ImageQuant LAS4000 (GE) or Odyssey XF (LICOR). The primary antibodies used were anti-FLAG-HRP (Sigma, 1:5000), anti-GFP (Roche, 1:2000), anti-actin (Proteintech, 1:5000), and anti-tubulin (Servicebio1, 1:5000).

For nuclear extraction, frozen seedlings (∼1.5 g per sample) were gently ground into powder with liquid nitrogen using a mortar and pestle. Nuclei were isolated with a protocol modified from Chen *et al.* ^43^. Briefly, ground powder was mixed in nuclei extraction buffer (Tris, Ficoll 400, Dextran T40, sucrose, MgCl_2_) with DTT, PMSF and cOmplete protease cocktail added right before use, then filtered through a 70 µm cell strainer followed by a 40 μm cell strainer. Filtered samples were adjusted with Triton X-100 to 0.5% (v/v) and incubated on ice for 15 mins. A small portion of sample was collected (total protein extract) before centrifugation at 1500 x g for 5 mins at 4 °C. After centrifugation, the supernatant (cytoplasmic fraction) was collected while the pellet (nuclear fraction) was washed three to four times by resuspending the pellet gently in nuclei extraction buffer with 0.1% Triton-X and then centrifuged at 1800 x g for 5 mins at 4 °C. The nuclear fraction was prepared by resuspending the pellet in 150 μL nuclei extraction buffer. Each fraction was mixed with 5X loading buffer and boiled at 95 °C for 10 mins for immunoblotting.

### Cycloheximide treatment

25 mM cycloheximide (CHX) stock (Sigma 01810) was prepared in Milli-Q water. For treatment, 7-day-old seedlings were submerged in 6-well plates containing 500 μM CHX in half-strength Murashige and Skoog liquid media. Samples were collected at indicated time points by drying briefly on Kimwipes and immediately frozen in liquid nitrogen to prevent further degradation.

### Heat treatment

For heat treatment, 10-day-old seedlings were treated directly on plates in Conviron growth chamber at 37 °C. Samples were collected at indicated time points and immediately frozen in liquid nitrogen to prevent further degradation. For confocal microscopy, 6-day-old seedlings were treated at 37 °C at indicated time points and imaged immediately after heat treatment.

### Confocal microscopy

To determine transient protein expression, Agrobacterium infiltrated *Nicotina benthamiana* leaf sections were imaged with Leica Stellaris 5 confocal microscope. To determine *in vivo* protein localization, root tips of 7-day-old *Arabidopsis* transgenic seedlings expressing CMT2 or CMT3 fused with GFP were imaged with Leica Stellaris 5 confocal microscope. To determine protein expression and localization after heat treatment, 6-day-old seedlings were imaged immediately after heat treatment with Leica SP8 Upright confocal microscope.

### Quantitative real-time PCR analysis

For RT-qPCR, plant total RNA was extracted using Ambion PureLink RNA Mini Kit (Invitrogen). First strand cDNA was then synthesized from 1 μg of the extracted total RNA using random hexamer, anchored oligodT_18_VN, murine RNase inhibitor and ProtoScript II (NEB) reverse transcriptase. For McrBC assay, plant genomic DNA was extracted with the CTAB method. An equal amount of genomic DNA was then digested with the methylation-dependent restriction endonuclease McrBC (NEB) for 8 hours at 37°C. The quantitative real-time PCR was performed in triplicates using iTaq Universal SYBR Green Supermix and ran with the Bio-Rad CFX Opus 96 Real-Time PCR System (Bio-Rad). The gene expression levels in RT-qPCR were normalized against wild type control and internal control *ACT7*. The relative methylation levels of McrBC-qPCR were normalized to uncut control.

### Protein expression and purification

Expression and purification of WT and mutant CMT2 follow a similar protocol as previously described for CMT3 ^21^. In short, each of the CMT2-encoding sequences was cloned into a modified PRSFDuet-1 vector (Novagen), fused with an N-terminal His6-SUMO tag. The expression plasmids were then transformed into *Escherichia coli* BL21 DE3 (RIL) cells for protein expression. The His6-SUMO-CMT2 proteins were purified through nickel affinity chromatography, followed by removal of His6-SUMO tag via ubiquitin-like-specific protease 1 (ULP1)-mediated cleavage. The tag-free CMT2 proteins were further purified through ion-exchange chromatography on a Heparin column (GE Healthcare) and size-exclusion chromatography on a 16/600 Superdex 200 pg column (GE Healthcare). The purified protein sample was stored in buffer containing 20 mM Tris-HCl (pH 8.0), 100 mM NaCl, 5 mM DTT and 5% Glycerol in a -80°C freezer before use.

### *In vitro* methylation assay

*In vitro* methylation assay was performed as described previously ^21^. In essence, The H3_1-17_K9me2 and H3_1-32_K9me2 peptides, each followed by a C-terminal tyrosine, was chemically synthesized, and verified by mass spectroscopy. Each 20-µL reaction mixture contains 0.5 µM wild-type or mutant CMT2 2 µM histone peptide, 2 µM synthesized DNA duplexes (hmCHG: 5′-AATATATXTGXAGXTGAATXAGXAGXTGTAATTTAA-3′, annealed with unmethylated, complimentary strand; X = 5mC; CHG: 5′- TGCTGCTGCTGCTGCTGCTGCTGCTGCTGCTGCTGCTGCTGCTGC-3′, annealed with unmethylated, complimentary strand; CHH: 5′- TACTACTACTACTACTACTACTACTACTACTACTACTACTACTAC-3′, annealed with complimentary strand), 0.56 µM S-adenosyl-L-[methyl-3H] methionine with a specific activity of 18 Ci/mmol (PerkinElmer), 1.96 µM nonradioactive SAM, 50 mM Tris-HCl (pH 8.0), 0.05% β-mercaptoethanol, 5% glycerol and 200 µg/mL BSA. The reaction lasted 30 minutes at 37°C before being quenched by addition of 5 µL of 10 mM cold SAM. Subsequently, 8 μL of each mixture was loaded onto Hybond N nylon membrane (GE Healthcare), which was left being dried out at room temperature. The membrane was then washed with 0.2 M ammonium bicarbonate (pH 8.2) three times, 5 minutes each, deionized water (5 minutes) once, and 95% ethanol (5 minutes) once. After air drying, the membrane carrying each sample was transferred into a vial containing 4 mL scintillation buffer (Fisher). The tritium scintillation was measured and recorded by a Beckman LS6500 counter. Each of the reactions was repeated three times.

### Bisulfite sequencing and data analysis

For whole genome bisulfite sequencing, seeds of Col-0, *cc*, and two lines of each indicated gCMT2-FLAG and corresponding V1200R mutation were planted on ½ MS medium for 10 days. Genomic DNA was then extracted from whole seedlings using the CTAB method. The genomic DNA was fragmented to a mean size of 100-300 bp by sonication using a Covaris S220 focused-ultrasonicator (Covaris), followed by end-repair, 3’-end adenylation and methylated adaptor ligation using Illumina TruSeq DNA kit (Illumina). Then bisulfite conversion was performed using a Zymo EZ DNA Methylation-Lightning kit (Zymo Research). The bisulfite-converted, adaptor-ligated DNA was enriched by PCR for 12 cycles using KAPA HiFi HotStart Uracil+ Kit (KAPA Biosystems), purified with Agencourt beads (Agilent) and quantified by Qubit HS dsDNA kit (Life Technologies). The integrity of the sequencing library was tested by Agilent 2100. The libraries were sequenced by 50 bp single-end method on a HiSeq4000 platform at NUcore sequencing center in Northwestern University (Chicago, IL, USA). Sequencing reads were trimmed using FASTP and aligned to the *Arabidopsis thaliana* TAIR10 genome using BSMAP version 2.9 allowing for 4% mismatches, trimming anything with a quality score of 33 or less, and removing any reads with more than five N’s ^44,45^. Methylation at every cytosine was determined by using BSMAP’s methratio.py script, processing only unique reads and removing duplicates. Differentially methylated regions (DMRs) were identified using methylKit with the following cutoffs for each sequence context: 40% difference for CG, 20% for CHG, and 10% difference for CHH DMRs and a p-value < 0.01 ^46^. Overlapping DMRs were identified using BEDTools *intersectBED* ^47^. TE metaplots were created using deepTools computeMatrix using BSMAP methylation file and a list of all TEs from TAIR10 ^48^. Whole genome methylation metaplots were created using the BSMAP methylation file and custom python scripts. Box plots were created using custom R scripts.

### RNA sequencing and analysis

For RNA sequencing, seeds of Col-0, *cc*, and two lines of each indicated gCMT2-FLAG and corresponding V1200R mutation were planted on ½ MS medium containing 1% sucrose for 10 days in long-day growth conditions (16 hrs light: 8 hours dark) at 22°C.Total RNA was extracted using PureLink RNA Mini Kit and treated with Dnase I. Library was constructed using a TruSeq RNA Library Preparation Kit (Illumina); messenger RNA was purified using RNA purification beads and fragmented by sonication using a Covaris S220 focused-ultrasonicator. One microgram of total RNA was reverse-transcried into cDNA with ProtoScript II. End-repair, 3’- end adenylation, ligation of adaptors and PCR amplification for 13 cycles were then performed. Reads were filtered using fastp and then aligned them to the TAIR10 genome using HISAT2 (v.2.0.0-beta) ^49^. Transcripts were assembled using StringTie^50^. The quantification of gene expression and the identification of DEGs were performed with DESeq2^51^. The heat map was made using Heatmapper (http://www.heatmapper.ca/expression/). Differentially upregulated and downregulated genes were analyzed using GO Term Enrichment tool on TAIR/PANTHER and fold enrichment was calculated by dividing the number of DEGs by the expected number of DEGs for a GO term ^52^.

### Other sequence analysis

For ChIP-seq analysis, raw data was downloaded for H3K9me2 (GSE111609) and H3K27me3 (GSE84483) were trimmed using Trimmomatic and aligned to the TAIR10 genome using Bowtie2 with the following parameter: -k 10 --very-sensitive --no-unal --no-mixed --no- discordant ^53,54^. For ATAC-seq analysis, raw data was downloaded from GSE85203 and aligned to the TAIR10 genome using Bowtie2 with the following parameter: -v 2 -m 3 ^54^. PCR duplicates were then removed using SAMtools ^55^. The uniquely mapped reads were kept and used for ChIP peaks calling with MACS2 ^56^. DMRs were overlapped with ChIP peaks using BEDTools *intersectBed ^47^*. Coverage bed graphs were generated using bamCompare for histone modifications normalizing to H3 and bamCoverage for ATAC-seq normalized with RPKM values ^48^. The level of each modification within the DMRs was calculated using a custom python script and box plots were generated using a custom R script.

### Sequence from multiple species and analysis

Sequences of CMT2 and CMT3 from multiple plant species were retrieved by BLAST using Ensembl Plants (plants.ensembl.org), NCBI, OneKP (db.cngb.org/onekp), PLAZA Gymnosperms (bioinformatics.psb.ugent.be/plaza/versions/gymno-plaza), FernBase (fernbase.org), or PhycoCosm (phycocosm.jgi.doe.gov). The sequences are listed in Supplementary Dataset 1. The disorder scores were determined using the VSL2 algorithm of PONDR (http://www.pondr.com/). The natural variations of CMT2 were from Araport11/TAIR JBrowse (https://jbrowse.arabidopsis.org/).

### Lhasa-0 sequence analysis

Paired end whole genome DNA sequencing libraries (utilized for SNV calling) were prepared by Starbio utilizing in house library preparation protocols. Paired end bisulfite sequencing libraries were prepared. 150 bp paired end sequencing was performed for both library types on an Illumina Hi-Seq 2500 instrument. Read alignment statistics can be found in Supplementary Table 1. To generate pseudoreference genomes for Tibet-0 and Lhasa-0, raw fastqs were croptrimmed for phred score >=30 using Trimmomatic and aligned to the TAIR10 genome (Col-0 ecotype) using Bowtie2 using the –local alignment setting, 1 bp mismatch in 22 bp seeds ^53,54^. SNVs and indels for Tibet-0 and Lhasa-0 were called against TAIR10 using the SAMtools mpileup function to compute likelihood of the data given each possible genotype and bcftools applied the priors to call variant sites. Annovar was used to do a gene based functional annotation of genetic variants ^57^. The variant called format (.vcf) files were used to produce a pseudoreference genome via the pseudoRef R-package (https://github.com/yangjl/pseudoRef).

For Tibet-0 and Lhasa-0 BS-seq analyses, data were initially aligned to the assembled pseudoreference genome via BSMAP version 2.9 ^45^. Reads were filtered for < 5 N, quality trimmed (Phred score >= 30) and alignment was allowed proceed with up to 8% read mismatch (default). Methylation tracks were created by calling cytosine methylation in all contexts via the methratio.py script removing duplicated reads and including zeros. Whole genome plots were produced by averaging DNA methylation in each context (CG, CHG, CHH) in 1 Mb bins via a custom python script (https://github.com/Sandman2127/Whole_Genome_DNA_Methylation_Plotter/blob/master/WG_methylation_plotter.py) and plotted using the R packages ggplot2 and reshape2 ^58,59^. After alignment via BSMAP, 100 bp DMRs were called against 54 wildtype high confidence Arabidopsis libraries via hcDMR caller, accepting only unique mappings and using the -Steve filter ^32,45^. Metaplots of DNA methylation were produced using deepTools compute Matrix and profiler functions at DMRs ^48^.

### 1001 Epigenome BS-seq data reanalysis

Data from the 1001 epigenomes project was reanalyzed using the above paramaters in BSMAP with previously built pseudoreference genomes (http://1001genomes.org/data/GMI-MPI/releases/v3.1/pseudogenomes/fasta/) ^60^. CHH methylation averages were averaged genome-wide using an awk script and plotted via the R-package rworldmap (https://CRAN.R-project.org/package=rworldmap).

### *dN*/*dS* analysis

Analysis of evolutionary constraint on the CMT2 gene by dN/dS was performed by generating a multiple sequence alignment for the CDS of CMT2 with *A. thaliana*, *A. lyrata*, *B. grandis*, *B. rapa*, *C. rubella*, and *B. distachyon* using ClustalW from Mega7 ^61^. Selection at each codon was estimated using the HyPhy ^62^. The data was interpreted according to the principles found previously ^63^.

### Statistical Analysis

The two-tailed Student’s t-tests were performed to compare distributions between different groups. For all tests, a p-value lower than 0.05 was statistically significant.

## Acknowledgements

We thank Ingrid Jordon-Thaden from Botany Garden and Greenhouses of University of Wisconsin-Madison for providing *Amborella trichopoda* materials. We thank Hiroshi Kawai (Research Center for Inland Seas of Kobe University, Japan) for providing *Chara braunii* plant material. This is in memory of Prof. Yang Zhong (Fudan University, ceased) for his work on collecting Arabidopsis in Tibet and providing Lhasa-0 seeds.

## Author contributions

JJ, JG, JF, and XJ performed the experiments. SML analyzed the high throughput sequencing data. DS analyzed the Lhasa-0 DNA methylation data. JJ, JS, and XZ conceived and organized the study. JJ, JG, and XZ prepared the manuscript with comments from all co-authors.

## Data and materials availability

Bisulfite-sequencing and RNA sequencing data have been deposited in NCBI Gene Expression Omnibus under accession number GSE247354. Seeds of Arabidopsis accessions and transgenic lines and plasmids generated in this study will be available upon request. Further information and requests for resources and reagents should be directed to and will be fulfilled by the lead contact, Xuehua Zhong.

## Supplementary Figures

**Figure S1.**
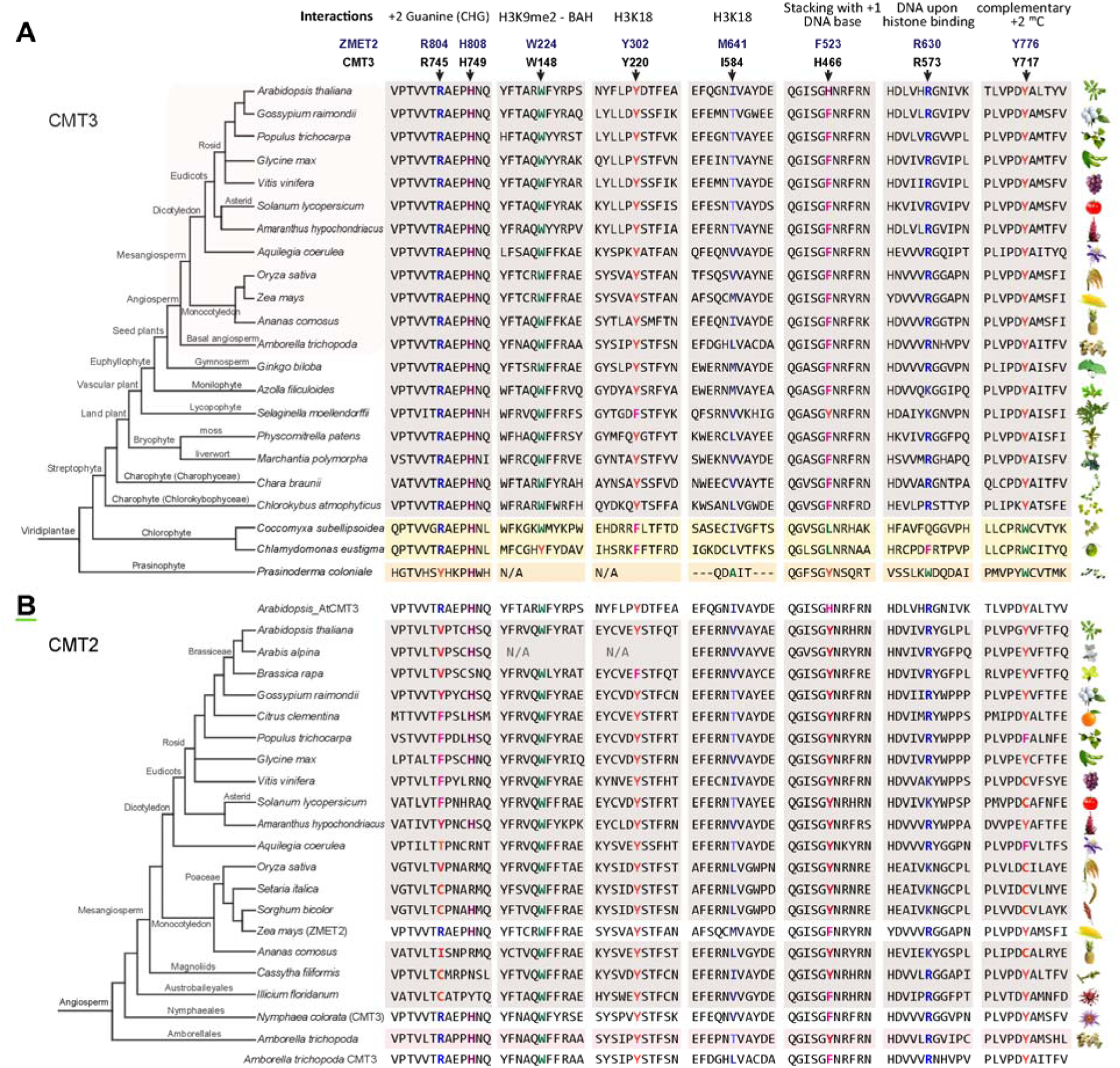
Key residues for CMT-type DNA methyltransferases in phylogenetic tree, related to Figure 1. (A and B) Phylogenetic tree depicting the conservation of key residues of CMT3 (A) and CMT2 (B) in plants. CMT3 is present in green plants *(Viridiplanlae),* while CMT2 is only present in flowering plants (angiosperms) and lost in some species during evolution.

**Figure S2.**
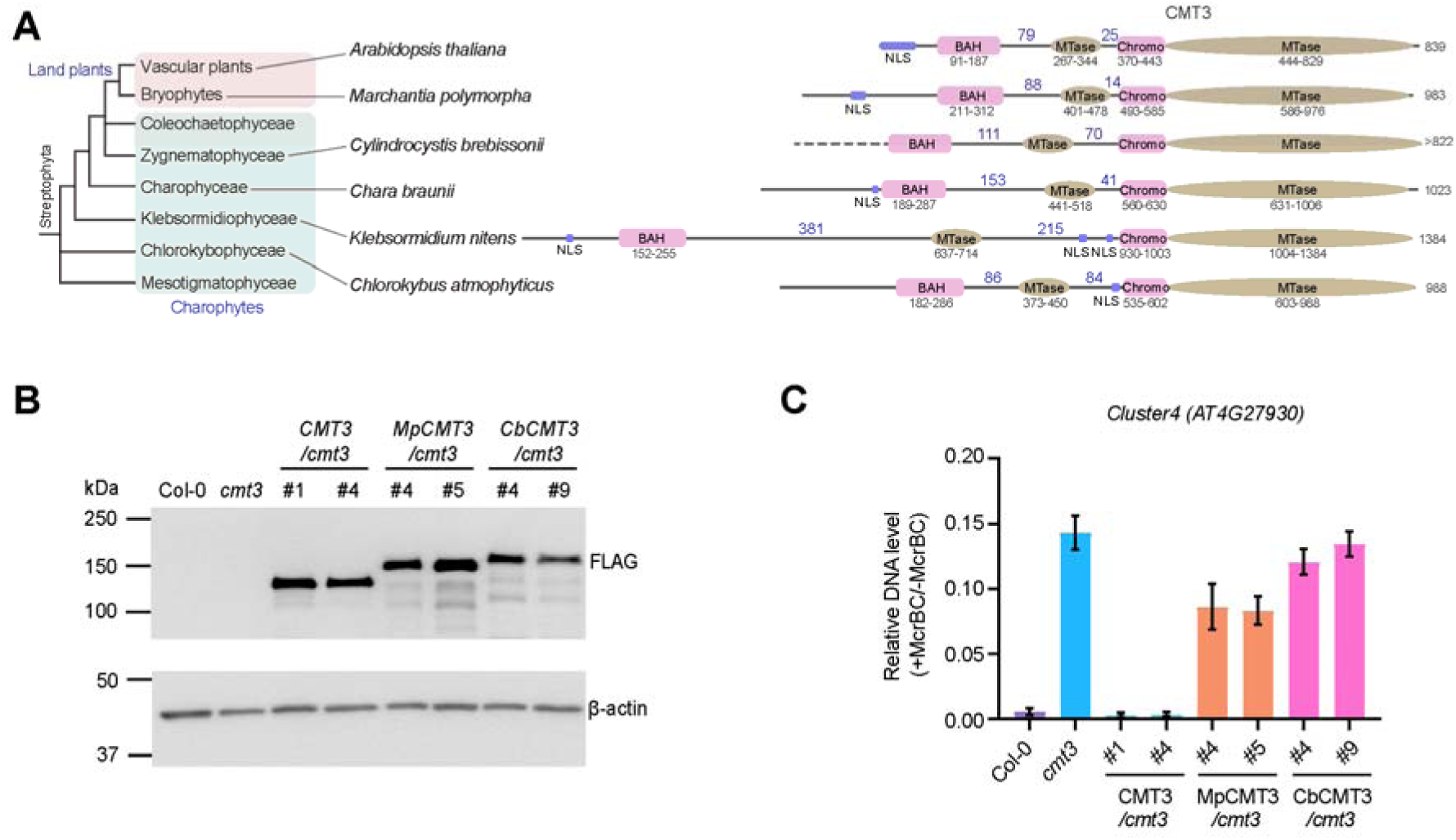
The CHG methylation activity of CMT3 is conserved across plant evolution, related to Figure 1. (A) Phylogenetic tree showing Charophytes and land plant species that share similar domain layouts in their CMT3 proteins. Nuclear localization signal (NLS) colored in violet. Numbers above lines depict the number of amimo acids between each domain. (B) Immunoblots showing protein levels of C. *braunii* and *M. polymorpha* CMT3 transformed into *Arabidopsis cmt3* mutant background under the control of UBQ10 promoter. (C) DNA methylation level over a CMT3 targeted locus, *Cluster*4, by C. *braunii* CMT3 (CbCMT3) and M. *polymorpha* CMT3 (MpCMT3) compared to wildtype (Col-0) and transgenic CMT3 measured by McrBC-qPCR assay.

**Figure S3.**
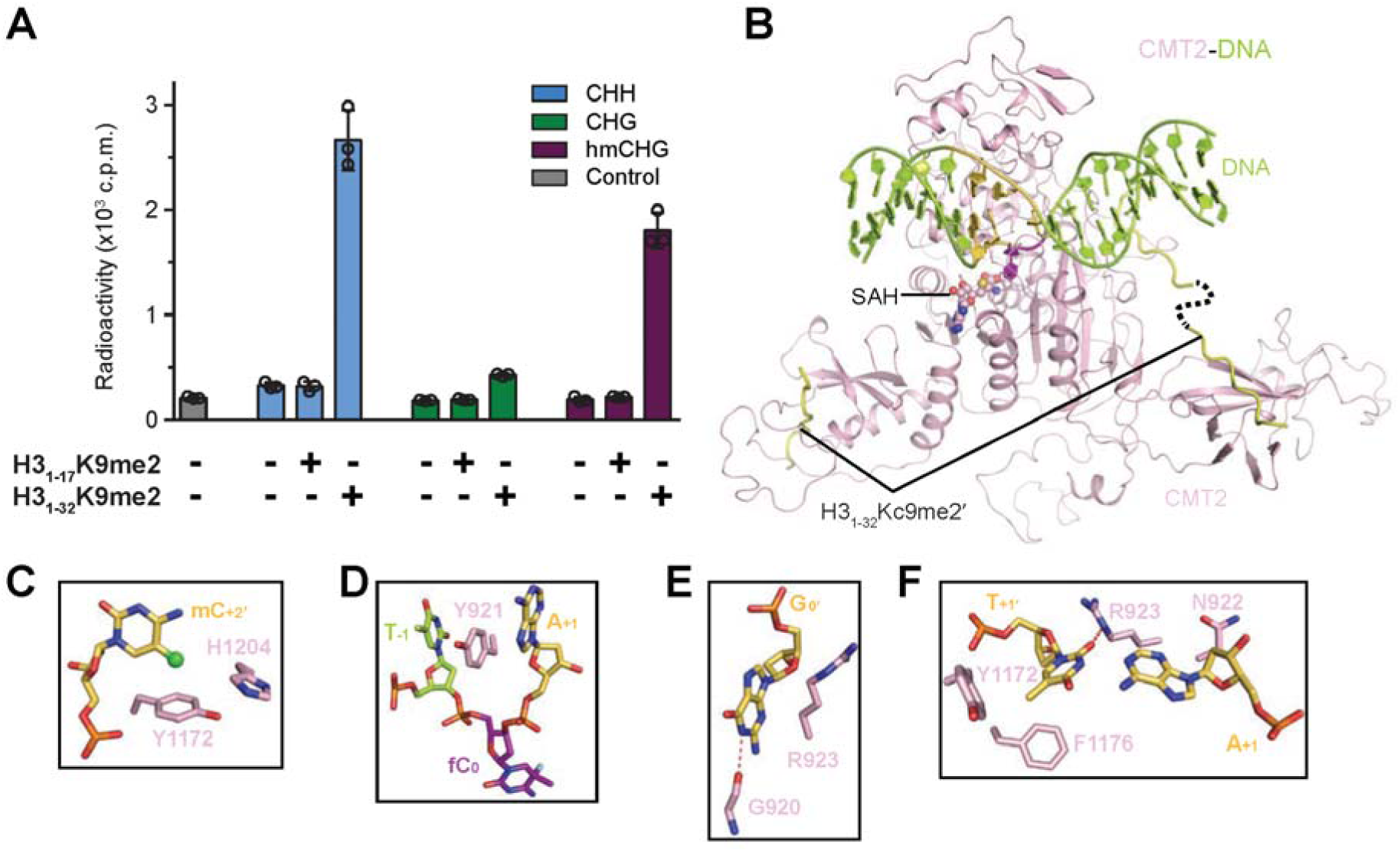
Predicted model of CMT2 interaction with DNA in CHG context, related to Figure 1. (A) *In vitro* DNA methylation assay of CMT2 on DNA duplex containing multiple CHH, CHG, or hemi-methylated CHG (hmCHG) sites, in the presence or absence of H3K9me3 peptides. Control, no substrate present. (B) Structural prediction of CMT2 interacting with DNA based on crystal structure of ZMET2^1^. (C) Close-up predictions of CMT2-DNA interactions at the +2-flanking mC_+2_. (D) Of CMT2-DNA interactions at the DNA cavity vacated by base flipping of fC_0_ on the target strand. (E) Of CMT2-DNA interactions at the orphan Go-site on the template strand (F) Of CMT2-DNA interactions at the +l-flanking A_+1_ T_+1_ pair.

**Figure S4.**
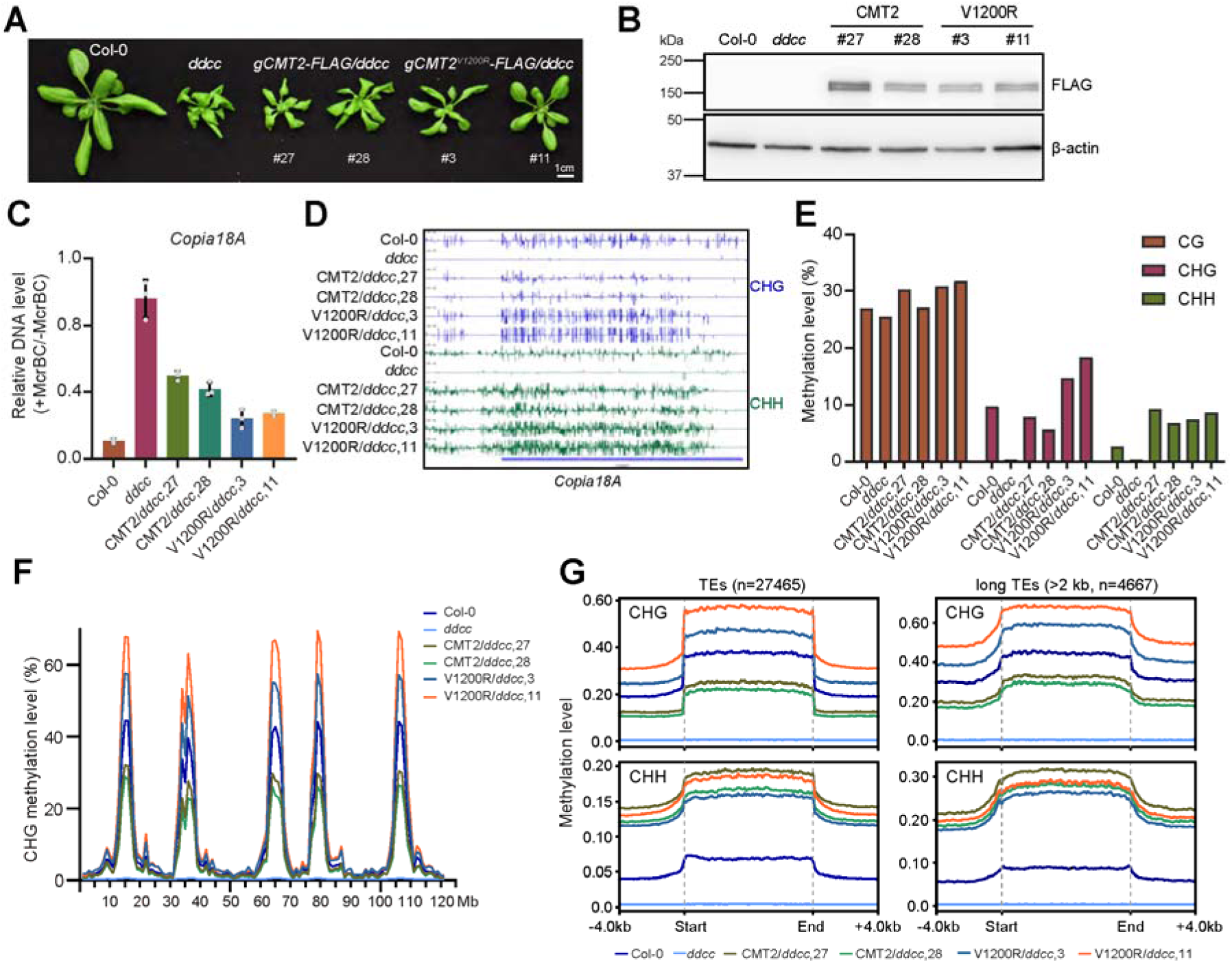
Effects of V1200R mutation on CHG methylation in *ddcc* mutant backgrounds, related to Figure 1. (A) Phenotypes of 3-week-old transgenic CMT2 and CMT2’^V1200R^ plants in *drmIdrm2cmt2cmt3* (*ddcc*) quadruple mutant background. (B) Immunoblots showing CMT2 and V1200R protein levels in *ddcc* background with actin as a loading control. (C) DNA methylation levels of CMT2 and V1200R over two CMT2-targeted TEs in *ddcc* measured by McrBC-qPCR assay and normalized with no-enzyme controls. (D) Genome browser view of CHG and CHH methylation levels of CMT2 and V1200R in *ddcc* background at *Copial8A* site. (E) Proportion of DNA methylation in CG, CHG, and CHH contexts of transgenic CMT2 and V1200R in *ddcc* background. (F) Metaplots for the CHG methylation across the Arabidopsis genome in CMT2 and V1200R in *ddcc* background. (G) The peaks represent the centromere and near centromere regions of the five chromosomes. Metaplots of CHG and CHH methylation over all TEs (n=27465) in Arabidopsis genome in CMT2 and V1200R in *ddcc* background and over long TEs (> 2kb, n=4667).

**Figure S5.**
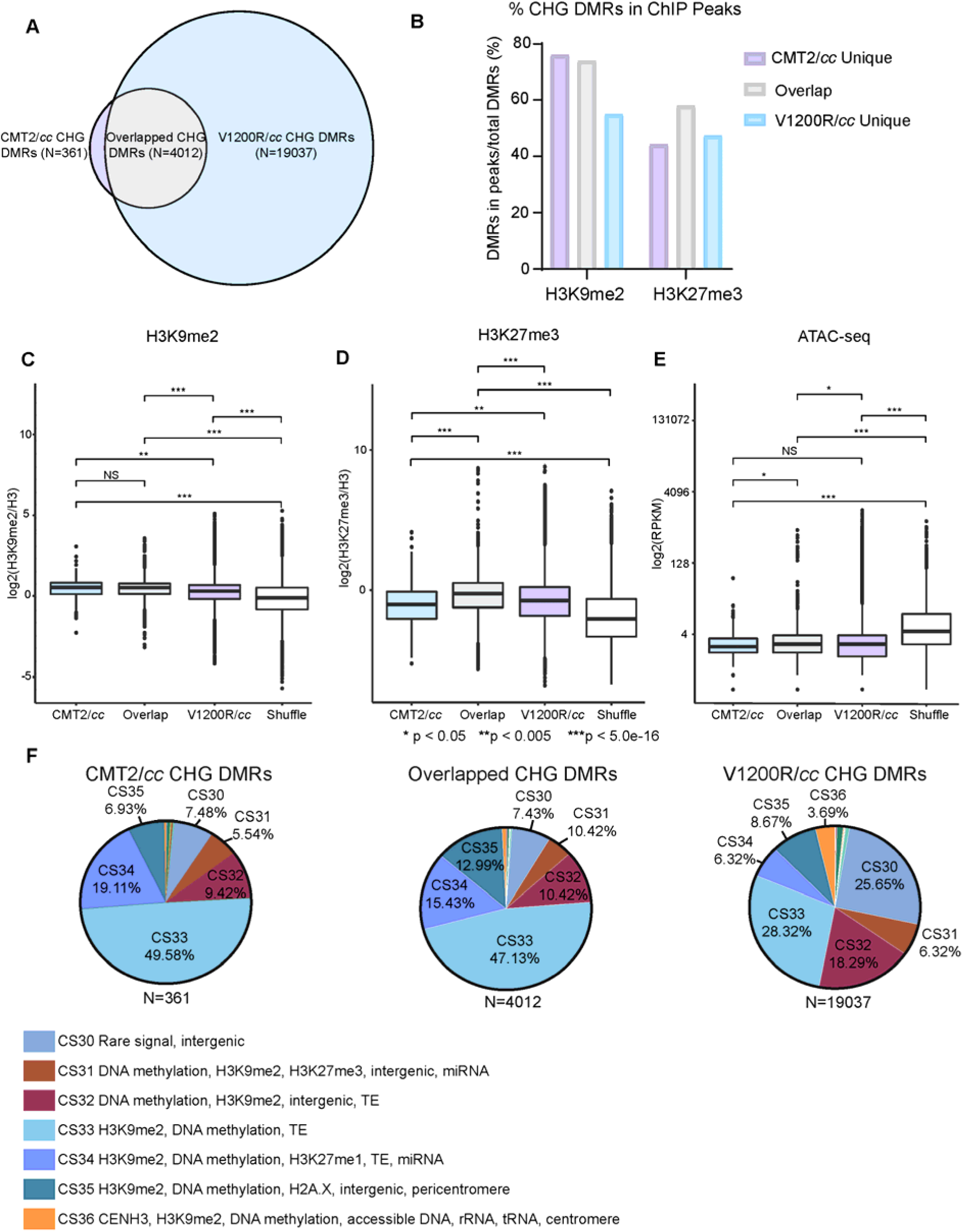
The chromatin environment of CMT2^Vl200R^ methylated regions, related to Figure 1. (A) Venn diagram depicting the overlap between CMT2 and V1200R CHG DMRs determined by the overlap in DMRs of two biological replicates. Only DMR demonstrated 100% overlap between two datasets is considered as an overlap. (B) Histogram showing the percentage of CHG DMRs shown in panel (A) that overlap with H3K9me2 and H3K27me3 ChIP peaks. Only DMR falls 100°% within the ChIP peaks is considered. (C-E) Boxplots showing the level of H3K9me2 (C), H3K27me3 (D), ATAC-seq (E) in Col-0. * p < 0.05, ** p < 0.005, ***p < 5. Oe-16 by Wilcoxon test. NS, not significant. (F) Pie charts showing the proportion of CHG DMRs falls into chromatin states defined by Liu *et al.^2^*.

**Figure S6.**
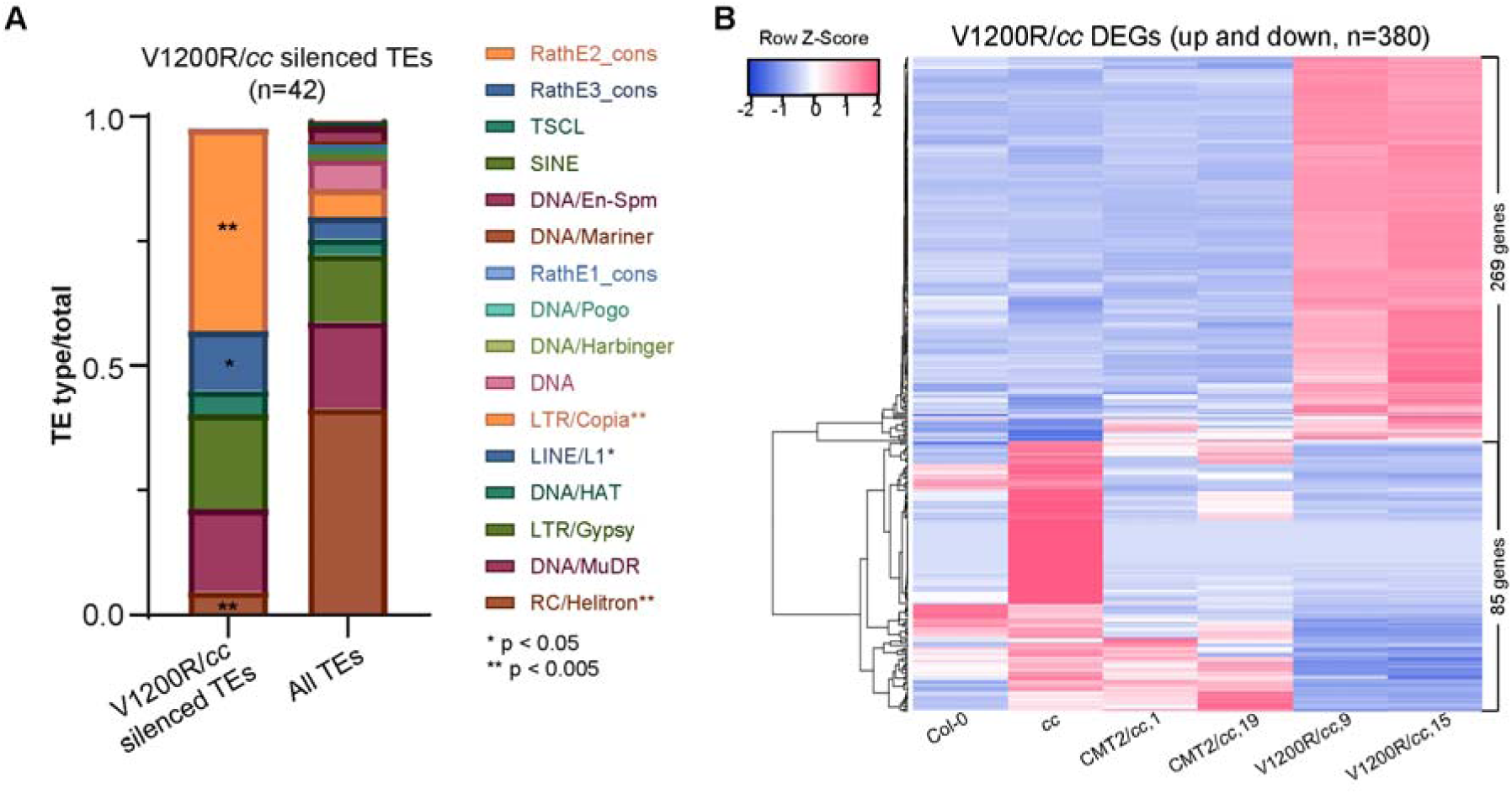
TEs and genes regulated by CMT2^Vl200R^, related to Figure 2. (A) Enrichment of TE types in TEs expressed in CMT2 but silenced in Col-0 and V1200R (shown in panel II). *p < 0.05, **p < 0.005 by Fisher’s Exact test. (B) Heat map showing the expression level of 380 shared DEGs (both up and down) between two biological replicates listed in Figure 2D.

**Figure S7.**
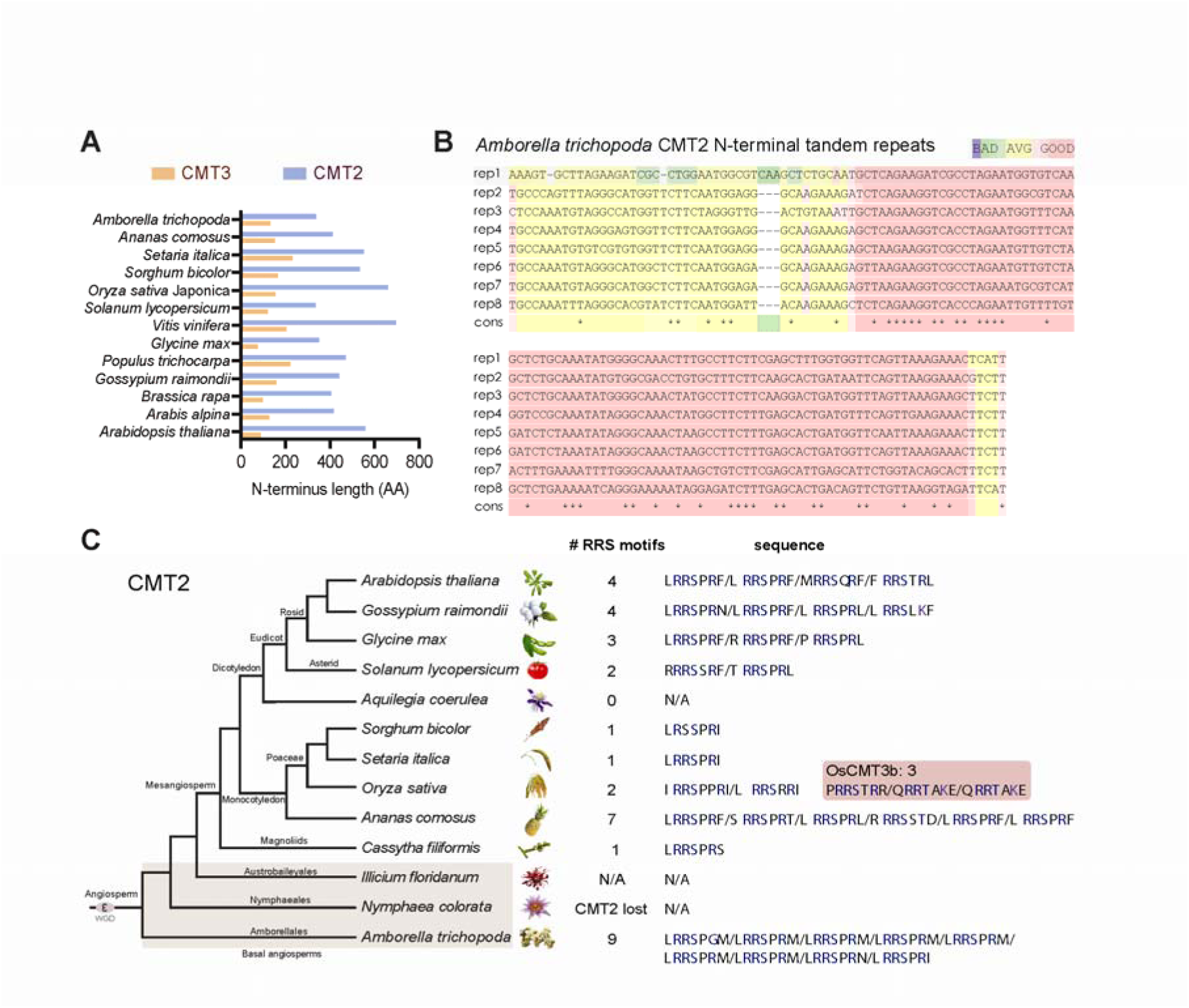
CMT2 RRS motifs are conserved in the angiosperm species, related to Figure 3. (A) N-terminus length of CMT3 and CMT2 from representative angiosperm species. (B) Alignment of the DNA sequences of eight tandem repeats in the N-terminus of *Amborella trichopoda* CMT2 by T-COFFEE. (C) The RRS motifs in CMT2 of representative angiosperm species.

**Figure S8.**
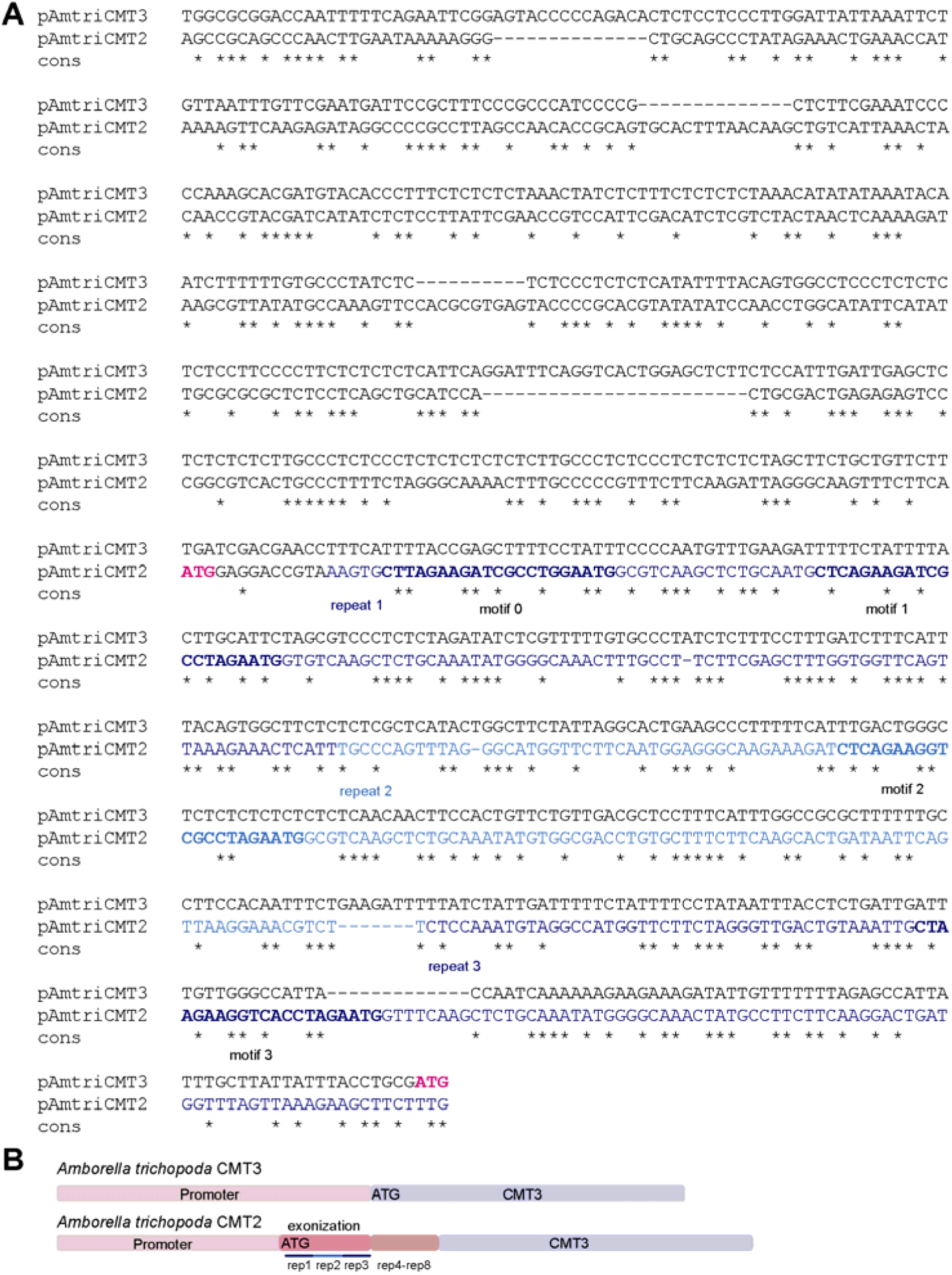
Alignment of *Amborella trichopoda* CMT2 and CMT3 promoter sequences, related to Figure 3. (A) Alignment of the promoter sequences of AmtriCMT3 and AmtriCMT2. (B) Diagram showing the gene structures of AmtriCMT3 and AmtriCMT2.

**Figure S9.**
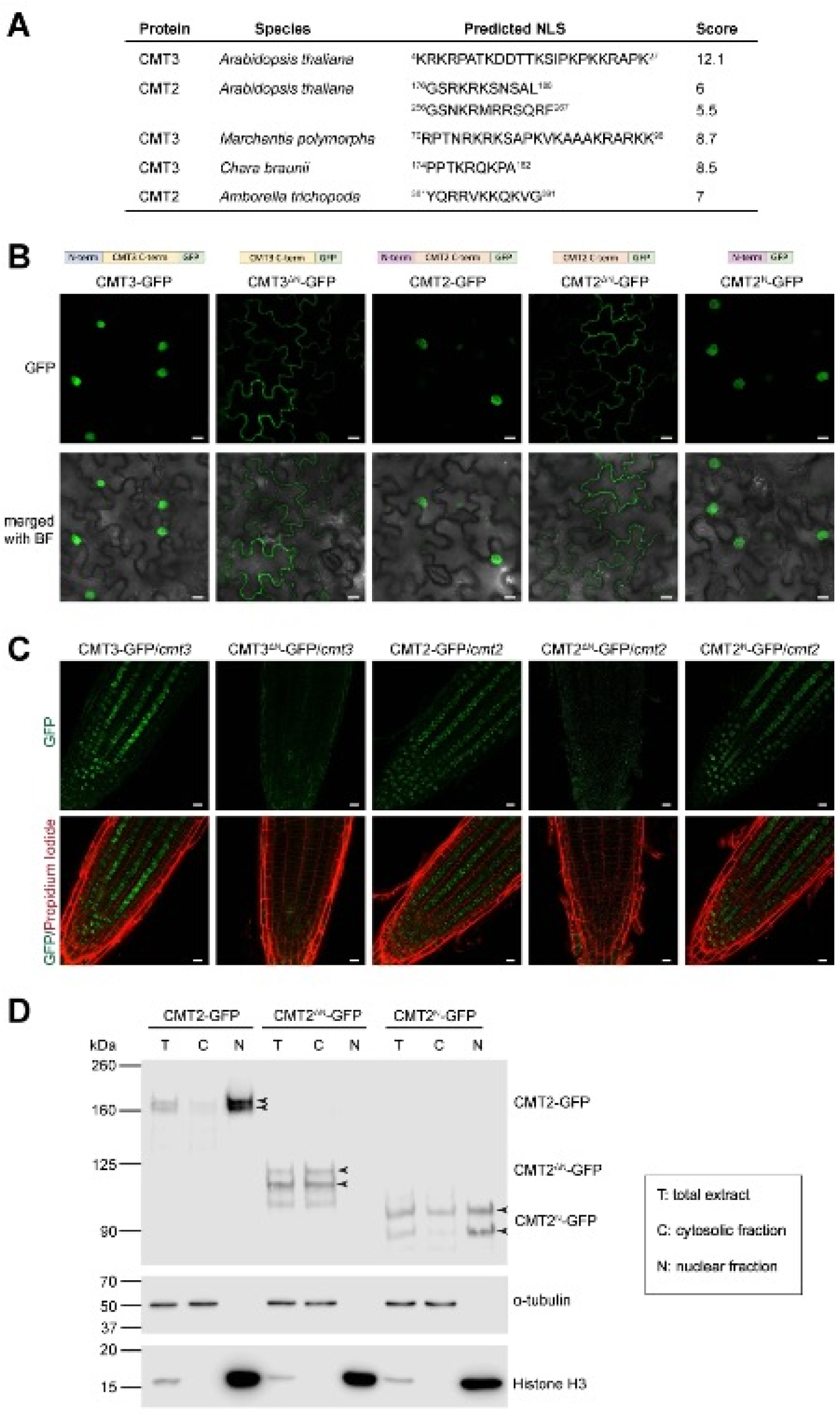
Nuclear localization signal dictates CMT2 and CMT3 localization, related to Figure 3. (A) Predicted nuclear localization sequence in CMT2 and CMT3 of different piant species by NLS mapper (https://nls-mapper.iab.keio.ac.jp). (B) Images Stowing GFP signals in *N. benthamiana* leaves transiently expressed with indicated CMT3 or CMT2 proteins (scale bar: 10μm) (C) Images showing GFP signals in *Arabidopsis* root tip of transgenic lines containing indicated CMT3 or CMT2 proteins (scale bar: 10μm). (D) Immunoblots detecting CMT2 protein levels in total (T), cytoplasmic (C), and nuclear (N) extracts from indicated transgenic plants.

**Figure S10.**
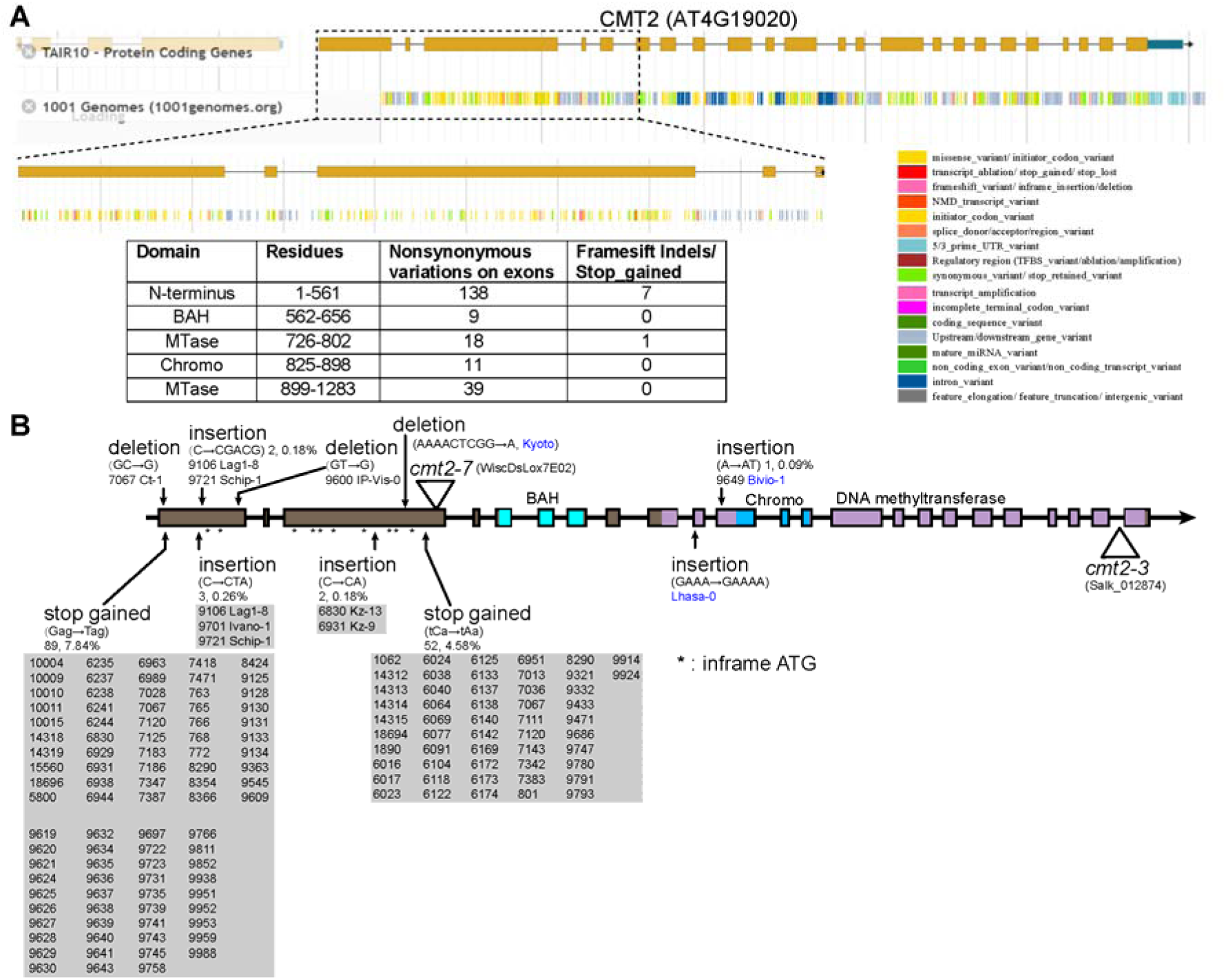
CMT2 natural variations from the 1001 genomes collection, related to Figure 6. (A) Browser snapshot of CMT2 natural variations and number of exon nonsynonymous variations on CMT2 domains. The data were from TA1R Jbrowse (https://jbrowse.arabidopsis.org). (B) Diagram showing the details of irameshiit-indels and stop-gained natural variations in CMT2. Kyoto is reported^3^. Lhasa-0 is from this study. * indicates possible novel inframe translation start sites.

**Figure S11.**
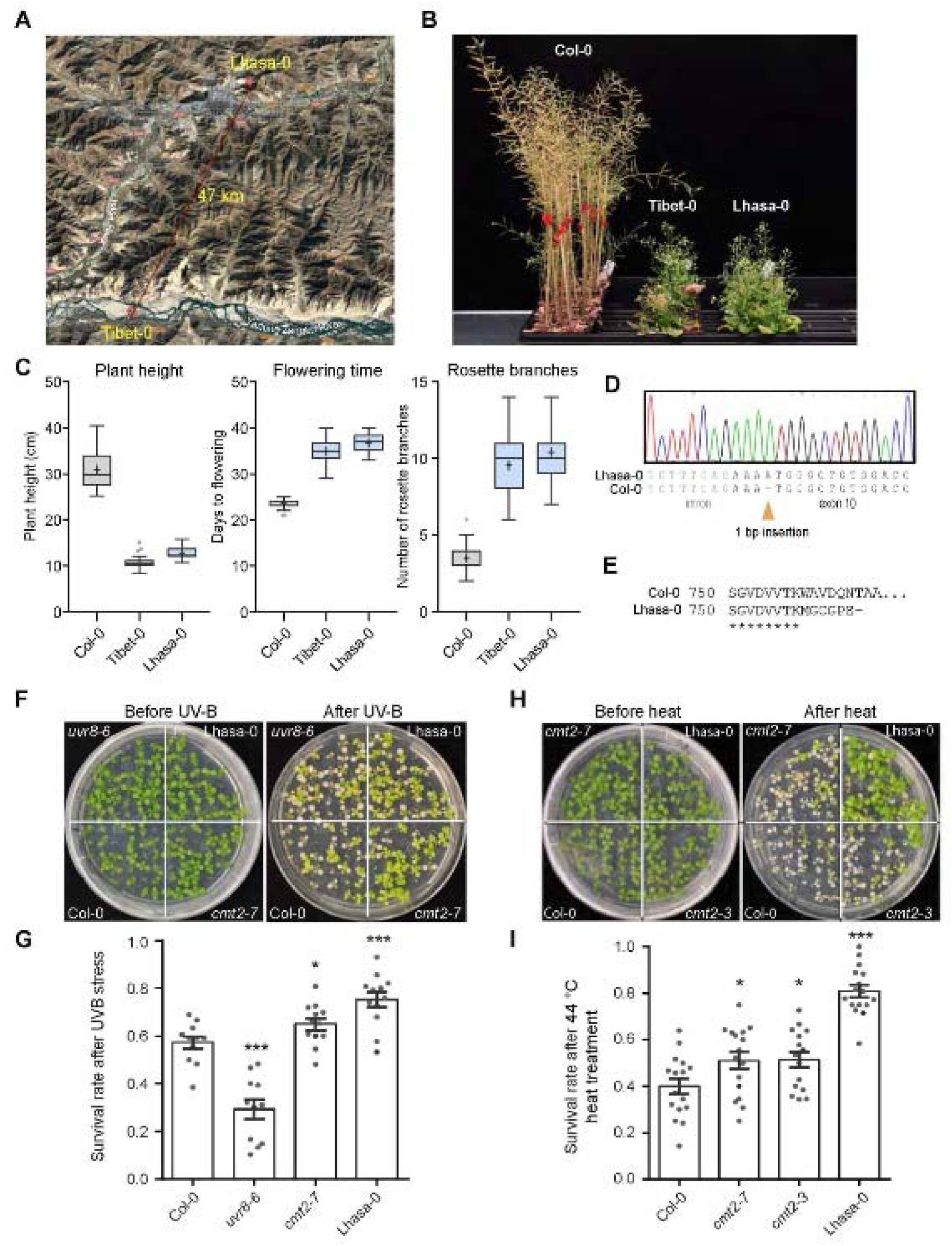
Characterization of *Arabidopsis thaliana* Lhasa-0 accession lacking functional CMT2, related to Figure 6. (A) Image of isolation region of Tibet-0^4^ and Lhasa-0 from the Tibetan Plateau near Lhasa. (B) Photos of Lhasa-0 at the flowering stage. (C) Phenotypes of plant height, flowering time, and rosette branches of Lhasa-0. Col-0 and Tibet-0 were used as controls. (D) Sanger sequencing conforming the 1 bp insertion in CMT2 of Lhasa-0. (E) Predicted amino acid sequence of the CMT2 mutation in Lhasa-0. (F) Images of seeding before and recovered from UV-B stress treatment. Treatment was performed for 3.5 hours on 10-day-old seedlings. (G) Survival rate in UV-B treatment from (F). Each grey dot represents a separate experimental replicate with n > 25 plants. (H) Images of seedings before and recovered after 2 hours basal heat treatment (44°C). Treatment was performed on 10-day-old seedlings. (I) Survival rate of plants in basal heat stress from (H). Each grey dot represents a separate experimental replicate with n > 25 plants.

**Figure S12.**
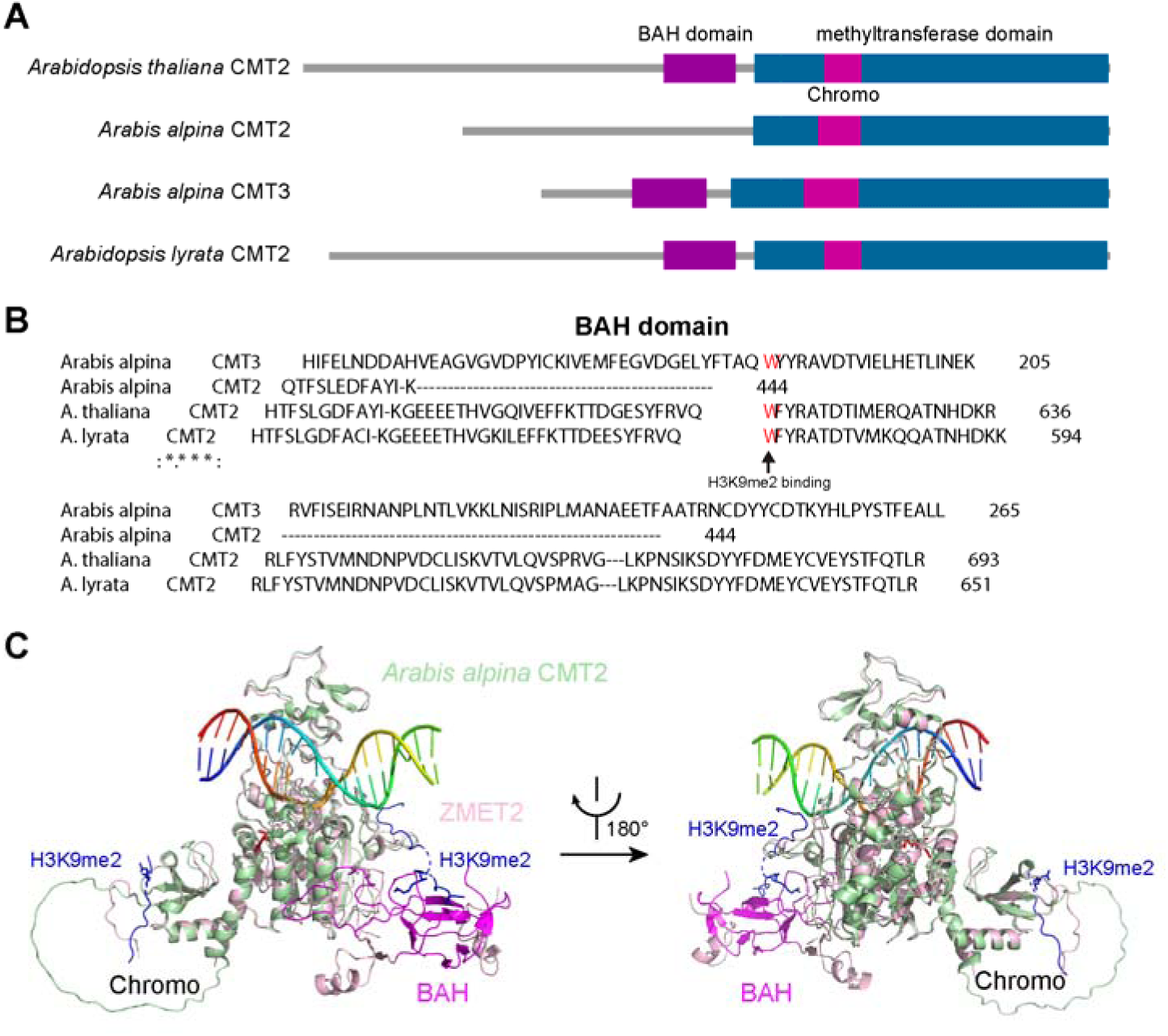
*Aracis alpina* CMT2 loses BAH domain, related to Figure 7. (A) Domains of CMT2 from *Arabidopsis thaliana* and *Arabis alpina*. (B) Alignment of amino acids showing the loss of BAH domain in *Arabis alpina* CMT2. (C) Predicted structure of Arabis alpina CMT2 by AlphaFold in comparison with a ZMET2-DNA-H3K9me2 Complex crystal structure^1^.

